# Tomato roots exhibit distinct, development-specific responses to bacterial-derived peptides

**DOI:** 10.1101/2024.11.04.621969

**Authors:** Rebecca Leuschen-Kohl, Robyn Roberts, Danielle M. Stevens, Ning Zhang, Silas Buchanan, Brooke Pilkey, Gitta Coaker, Anjali S. Iyer-Pascuzzi

## Abstract

- Plants possess cell-surface recognition receptors that detect molecular patterns from microbial invaders and initiate an immune response. Understanding the conservation of pattern-triggered immunity within different plant organs and across species is crucial to its sustainable and effective use in plant disease management but is currently unclear.
- We examined the activation and immune response patterns of three pattern recognition receptors (PRRs: *Sl*FLS2, *Sl*FLS3, and *Sl*CORE) in different developmental regions of roots and in leaves of multiple accessions of domesticated and wild tomato (*Solanum lycopersicum* and *S. pimpinellifolium*) using biochemical and genetic assays.
- Roots from different tomato accessions differed in the amplitude and dynamics of their immune response, but all exhibited developmental-specific PTI responses in which the root early differentiation zone was the most sensitive to molecular patterns. PRR signaling pathways also showed distinct but occasionally overlapping responses downstream of each immune receptor in tomato roots.
- These results reveal that each PRR initiates a unique PTI pathway and suggest that the specificity and complexity of tomato root immunity are tightly linked to the developmental stage, emphasizing the importance of spatial and temporal regulation in PTI.

## INTRODUCTION

Plants exhibit a multi-layered defense system, comprised of pre-formed barriers and induced defense responses. Constitutive defenses of the plant often include physical barriers such as cell walls, waxy epidermal cuticles, or targeted lignin deposition aimed to restrict pathogen movement (Malinovsky et al., 2014; Serrano et al., 2014; Kashyap et al, 2022). Defense responses can be induced by pathogen recognition, either through pattern-triggered immunity (PTI) or effector-triggered immunity (ETI) (Yuan et al., 2021; Yu et al., 2024). In PTI, cell surface-localized receptor proteins known as pattern-recognition receptors (PRRs) identify foreign signatures from a pathogen in the initial stages of invasion. These signatures, also called pathogen-associated molecular patterns (PAMPs) or microbe-associated molecular patterns (MAMPs), are found throughout a range of microbes, from soil-borne bacteria to foliar fungi (Miya et al., 2007; Wei et al., 2018; Luo et al., 2023). Upon recognition of a PAMP/MAMP, the host initiates defense responses including the short-term formation of reactive oxygen species, increased calcium signaling, activation of mitogen-activated protein kinase cascades, halted growth, and transcriptional reprogramming – altogether known as PTI (Shu et al., 2023).

MAMPs are well-conserved across microbial species, and many PRRs recognize more than one pathogen (Cheng et al., 2021; Colaianni et al., 2021; Ngou et al., 2022). Known MAMPs are highly conserved across pathogens, making it less likely for them to develop mutations that evade PRR recognition (Zhao et al., 2022). Interspecies transfer of PRRs can expand crop resistance (Frailie et al., 2021), and PRR-based crop engineering has the potential to provide broad-spectrum and durable resistance (Lacombe et al., 2010, Li et al. 2024). However, for this to be a sustainable strategy, detailed knowledge of PTI in crops is needed.

Key knowledge of PTI originates from work in *Arabidopsis thaliana* (Arabidopsis) and the well-characterized leucine-rich repeat receptor-like kinase (LRR-RLK) PRRs FLAGELLIN SENSING2 (*FLS2*) (Gomez-Gomez, 1999/2000; Chinchilla et al., 2006), and EF-Tu RECEPTOR (EFR) (Zipfel et al., 2006; Ngou et al., 2022). Recognition of flg22 by FLS2 or elf18 by EF-Tu activates a suite of downstream responses that includes a complex network of co-receptors, receptor-like cytoplasmic kinases (RLCKs), calcium-dependent protein kinases (CDPKs) and mitogen-activated protein kinases (MAPKs) (Asai et al., 2002; Boudsocq et al., 2010; Li et al., 2014; Lee et al. 2021). Both *At*FLS2 and *At*EFR-driven responses require the co-receptor *At*BAK1 (brassinosteroid insensitive 1-associated receptor kinase 1) and RLCK *At*BIK1 (Botrytis-induced kinase 1) for the initiation of ROS burst by *At*RBOHD and activation of the *At*MAPK signaling pathway.

In tomato (*S. lycopersicum*) three PRRs have been identified in bacterial-plant interactions: *Sl*FLS2, the receptor for flg22; FLAGELLIN SENSING3 (*Sl*FLS3), the receptor for flgII-28; and *Sl*CORE, the receptor for csp22 (Robatzek et al., 2007; Hind et al., 2016; Wang et al., 2016); a tomato EFR homolog is not found in the genome. While *FLS2* is present in most plant genomes, both *FLS3* and *CORE* are found in the Solanaceae family (Felix and Boller, 2003; Clarke et al., 2013; Wang et al., 2016). *Sl*FLS2 signaling in tomato has both some similarities and differences compared to Arabidopsis (Nguyen et al., 2010; Roberts et al., 2020). The tomato orthologs of *At*BAK1, *Sl*SERK3A/3B (somatic embryogenesis receptor kinase 3A/ 3B), interact with *Sl*FLS2 and trigger downstream ROS response and root growth inhibition (RGI) (Peng & Kaloshian, 2014). Interspecies transfer of *At*EFR to tomato resulted in resistance to the soil-borne bacterial pathogen *Ralstonia solanacearum* (Lacombe et al., 2010, Kunwar et al. 2018), suggesting that molecular components downstream of *At*EFR are conserved in tomato. However, it is unclear exactly which elements and molecular mechanisms of PTI are conserved.

A previous screen of heirloom tomatoes revealed natural variation in ROS response to all three MAMPs (flg22, flgII-28, and csp22), both between type of MAMP and within cultivars (Veluchamy et al, 2014; Roberts et al., 2020). This, along with variations in temporal dynamics of MAPK responses flg22 or csp22 perception in *N. benthamiana* transient assays (Wei et al., 2018), led us to hypothesize that although the basic tenets of PTI are conserved between FLS2, FLS3 and CORE, the details of the downstream molecular signaling vary among receptors.

Immune signaling pathways and their associated proteins have focused on foliar tissues, but plant roots can differ in their immune responses compared to above-ground counterparts and show targeted expression of PRRs within different tissue types (Beck et al., 2014; Chuberre et al., 2018). For example, *At*EFR is primarily found in aboveground, reproductive tissues and does not activate a ROS response upon recognition of elf18 in Arabidopsis roots (Wyrsch et al., 2015). Understanding how root tissues differ in their PTI response is imperative for implementation of PRR transfer for broad-spectrum resistance, as soil borne pathogens such as *Ralstonia solanacearum* enter through wounds or natural openings in the root tissues.

Here we investigate PTI signaling and responses in tomato roots, tracing the pathway from cell surface recognition to downstream phenotypic outcomes. Through characterization of PTI response in *S. lycopersicum* and *Solanum pimpinellifolium* roots to PAMPs flg22^Pst^, flgII-28^Pst^, and csp22^Rsol^, we reveal that ROS species formation, root growth inhibition, and intermediate signaling components vary both between PAMP treatment type and across various cultivars. We show that PTI response is primarily found in early differentiation regions of roots, including ROS burst and MAPK activation, underscoring the importance of these areas in early defense signaling.

Finally, we show that tomato root PTI responses vary from those in Arabidopsis, including a lack of seedling growth inhibition for flgII-28^Pst^ and csp22^Rsol^ treatments and differential regulation of ROS burst by *Sl*SERK3A/3B. Our results show that the molecular details of the signaling downstream of *Sl*FLS2, *Sl*FLS3, and *Sl*CORE differ, and highlight the need for further characterization of root PTI pathways.

## MATERIALS AND METHODS

### Plant Material and Plate Growth Conditions

Tomato accessions listed in **Table 1** were sterilized for 10 minutes in 50% bleach, then washed three times with water. Seeds were plated on 1% agar plates at 4° C overnight before placing at room temperature (22° C) at a 16:8 h day/night cycle.

**Table 1.**
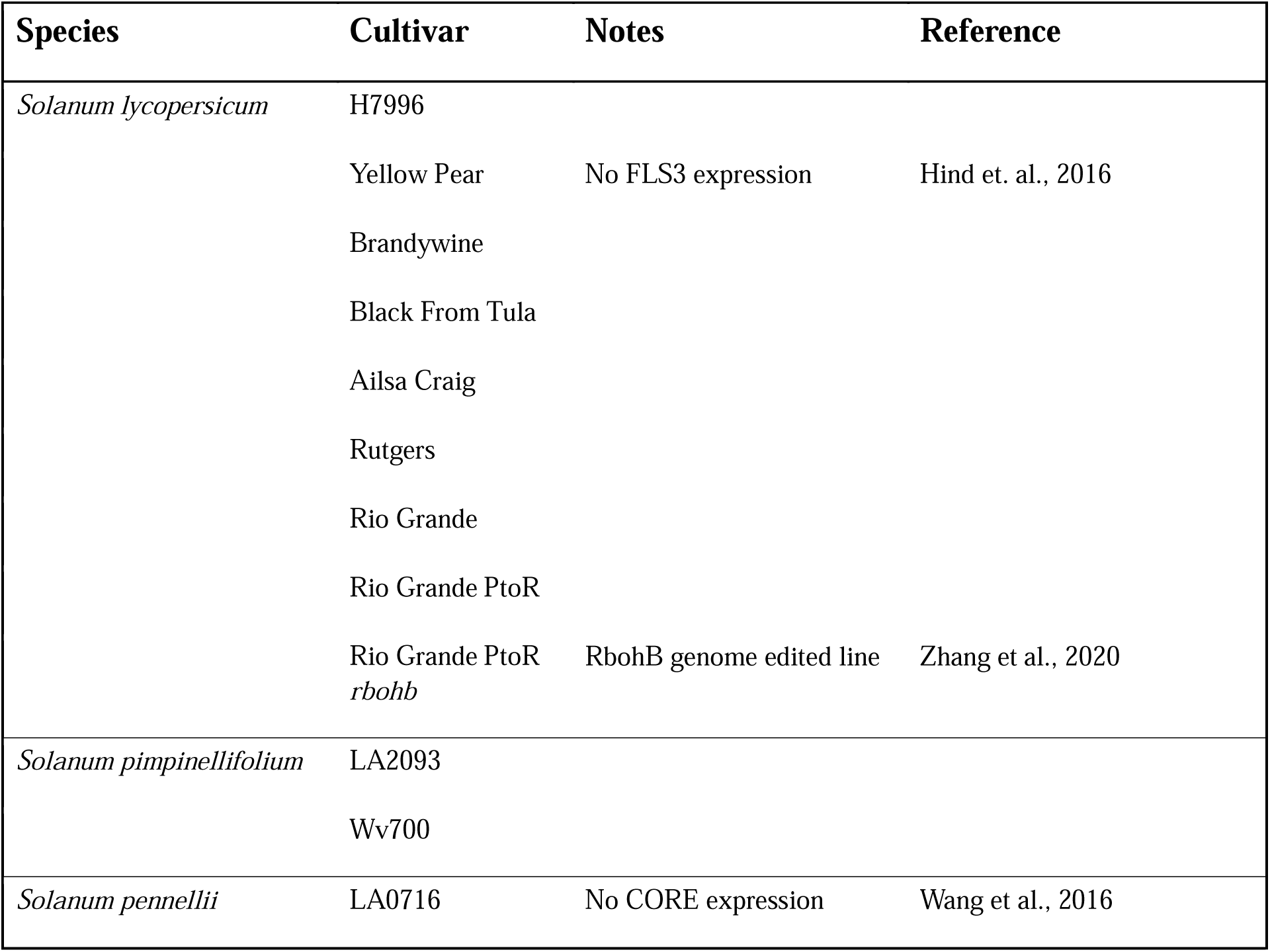
Cultivars of tomato used in this study.

*Arabidopsis thaliana* seeds (Col-0, *AtrbohD, AtrbohF, AtrbohD/AtrbohF*) were sterilized for 5 minutes in 50% bleach and 0.001% Tween, then washed three times with water. Seeds were stratified in ddH20, then covered for 48 hours at 4° C before plating on 0.5X Murashige and Skoog (MS) medium, 1% sucrose. Seeds were grown in a controlled chamber at 22° C at a 16:8 h day/night cycle. Mutant seeds were obtained from the lab of Chris Staiger, Purdue University Department of Botany and Plant Pathology.

### Generation of *rbohb* Mutant in Tomato

Mutant seeds (Rio Grande PtoR – *SlrbohB*) were generated using genome editing approaches as previously described in Zhang et al. (2020). To generate the *rbohb* mutant in the tomato (*Solanum lycopersicum*) cultivar Rio Grande (RG)- PtoR, one guide RNA (gRNA: 5’- GGACCGCTGAACAAACGAGG-3’) was designed to target the first exon of *RbohB* (Solyc03g117980). The gRNA cassette was cloned into the p201N:Cas9 binary vector and tomato transformation was performed at the Biotechnology Center at the Boyce Thompson Institute as described previously (Jacobs et al., 2015; Jacobs et al., 2017). The *rbohb* mutant line used in this study carries a 1 bp insertion in the first exon of the *SlRbohB* gene, resulting in a loss-of-function mutation in *SlRbohB* in the plants. Mutations were confirmed by PCR amplification using primers found in **Supplemental Table 1** and Sanger sequencing. Lines were verified to be homozygous, biallelic mutants and Cas9 was segregated out.

### Peptides

flg22^Pst^ and csp22 peptides were purchased from EZBiolabs, using the following amino acid sequences: flg22^Pst^ QRLSTGSRINSAKDDAAGLQIA; csp22^Rsol^: ATGTVKWFNETKGFGFITPDGG.

The flgII-28^Pst^ and flg22^Rsol^ peptide was purchased from GenScript, with the following amino acid sequence: flgII-28^Pst^: ESTNILQRMRELAVQSRNDSNSATDREA, flg22^Rsol^ QRLSTGLRVNSAQDDSAAYAAS.

### Temporary Root Growth Inhibition (RGI) Assay

Tomato seedlings were grown on 1% water agar plates in the conditions as described above. Four-day old seedlings were scanned and treated with 300 µL of elicitor treatment (1 µM flg22^Pst^, 100 nM flgII-28^Pst^, 1 µM csp22^Rsol^, or water), making sure to only submerge the root organ. Tomato seedlings were then scanned again at 24- and 48-hours post inoculation and measured using ImageJ for subsequent analysis.

Arabidopsis seedlings were grown on 0.5X MS, 1% sucrose in the conditions as described above. Five-day old seedlings were scanned and treated with 200 µL of elicitor treatment (1 µM flg22^Pst^ or water), making sure to only submerge the root organ. Arabidopsis seedlings were then scanned again at 24- and 48-hours post inoculation and measured using ImageJ for subsequent analysis.

### Oxidative Burst Luminescence Assay

The ROS assay was performed on tomato roots as described previously with a number of modifications (Wei et al., 2018). For whole-root assays, tomato seedlings were grown on 1% agar in the conditions described above. Five-day old tomato roots were placed under microscope and cut at the root-shoot junction. For developmental zone assays, the five-day old tomato roots were placed under a microscope and cut at the point of first visual root hair, the point at which root hairs were fully emerged, and at the root-shoot junction. The early differentiation zone (ED) was defined as the root section exhibiting emerging root hairs, while the late differentiation zone (LD) exhibited fully emerged root hairs. All root segments were then weighed with a precision balance before being placed in a white 96-well plate (Perkin Elmer, OptiPlate-96) with 200 µL of fresh water to recover. Segments were washed with water and kept in the dark for one hour, after which the water was removed, and fresh water was placed in each well and sat overnight in darkness. After overnight recovery, the water was removed and replaced with 200 µL of the corresponding master mix for each peptide elicitor. Master mix was made from 500X L-012 stock solution (LSS) and 500X horseradish peroxidase stock solution (HPSS) and the corresponding peptide for a final concentration of 1.5X L-012 (Wako Chemicals USA) and 1.5X HPSS (Thermo Fisher Scientific). Master mixes used had a final peptide concentration of 1 µM flg22^Pst^, 100 nM flgII-28^Pst^, or 1 µM csp22^Rsol^. Relative light units (RLUs) were detected using an Infinite 200 Pro Luminescent Microplate Reader (Tecan Life Sciences, Switzerland) and exported to an excel spreadsheet for further analysis. Three technical replicates were used for each analysis, with six roots per treatment. Data were normalized and expressed as RLU per milligram of fresh weight.

For tomato leaves, ROS assays were performed as previously described in (Hind et al., 2016) using 100 nM of DC3000 flg22 or flgII-28 peptides. The average ROS response for each plant is the mean of three replicate leaf discs from four plants. The assay was performed on ten independent VIGS biological replicates with similar results, and one representative experiment is shown in Figure 3.

### Cloning

Constructs used in the *in vitro* kinase assays were amplified via PCR using the primers found in **Supplemental Table 1**. Total RNA was extracted from tomato (Rio Grande) using the Qiagen RNeasy Plant Mini Kit (Cat. 74904) and used to generate cDNA (Invitrogen SuperScript III, 12574018). The cytoplasmic domains of the SERKs (SERK3A-CD and SERK3B-CD) were PCR-amplified from cDNA and inserted into the Gateway vector pDONR/Zeo (Invitrogen, 12535035) following the manufacturer’s instructions. Sequences were confirmed via Sanger sequencing, and then the construct ORFs were cloned into Gateway vector pDEST-HisMBP (Nallamsetty et al., 2005) using the LR Clonase II enzyme following the manufacturer’s instructions. Mutagenesis of the SERK3A-CD and SERK3B-CD clones was performed in the entry vectors using a Q5 Site-Directed Mutagenesis kit following the manufacturer’s instructions (New England Biolabs; www.neb.com).

### Virus-induced gene silencing (VIGS)

The pTRV vector derivatives (pTRV2-EC1, pTRV2-SlSERK3A, pTRV2-SlSERK3B, and pTRV2-SlSERK3A/3B) were transformed into *Agrobacterium tumefaciens* strain GV3101and prepared for infection (final OD=0.5) in tomato seedlings as previously described (del Pozo et al., 2004). Knockdown of gene expression was confirmed in qPCR using the primers in **Supplementary Table 1** as described previously (Mantelin et al., 2011). VIGS experiments were repeated a total of ten times using four plants per replicate (n=40 for each VIGS construct) with similar results.

### *In vitro* Kinase Assays

HisMBP-tagged proteins were expressed and induced in BL21 (DE3) pLys Rosetta cells as described previously (Roberts et al., 2020). *In vitro* kinase assays were performed for 30 minutes at room temperature in 20µL of reaction buffer (50mM HEPES, pH 7.5, 10mM MgCl_2_,10mM MnCl_2_, and 2 µCi [γ-^32^P]) using 5 μg of each of the various kinase proteins and/or 3 μg of myelin basic proteins, as previously described (Roberts et al., 2019). The assay was repeated a total of six times with similar results.

### Treatments of Diphenyleneiodonium Chloride (DPI)

To determine the concentration of diphenyleneiodonium chloride (Sigma Aldrich, CAS: 4673- 26-1) required to inhibit ROS burst caused by flg22^Pst^, the oxidative burst luminescence assay above was repeated with mock, 1 uM flg22^Pst^, and 1uM flg22^Pst^ solutions containing a final concentration of DPI between 0-1 uM.

Root growth assays including DPI were treated one hour before inoculation with 1 uM DPI as determined by the oxidative burst luminescence assay referenced above. The roots were then treated at 0 hpi with an elicitor solution of mock, 1 uM flg22^Pst^, or 1 uM flg22^Pst^ and1 uM DPI.

### Plant Growth of Tomato Accessions in Soil

H7996 (*S. lycopersicum*) was sterilized using the above method. Seeds were stratified in water and left at 4 C overnight before planting. Plants were grown in conditions as described in Meline et al. (2022) with slight modifications. Seeds were grown in BM3 in 3.8 cm x 8.6 cm x 5.8 cm (L x W x D) at 28°C and 16/8 h day/night. Twelve days after germination, plants were treated with 28 mL of Peter’s Excel Fertilizer (86.4g/L).

### Determination of MAPK Phosphorylation

Tomato (H7996) 5-day old seedlings were cut from the above-ground tissues at the root-shoot junction and further separated into whole root samples, late differentiation zone samples, or early differentiation zone samples. The root segments were allowed to sit for six hours in ddH_2_0 before being placed into a solution of 1 µM flg22^Pst^, 100 nM flgII-28^Pst^, or 1 µM csp22^Rsol^. The tissue was harvested at 0- or 10-minutes post treatment and flash frozen in liquid nitrogen. For tomato leaves, leaf discs were collected from eight-week-old tomato leaves (H7996) and allowed to sit for six hours before being placed into a solution of 1 µM flg22^Pst^, 100 nM flgII-28^Pst^, or 1 µM csp22^Rsol^ and flash frozen in liquid nitrogen after 10 minutes.

Total proteins were extracted using a protein extraction buffer (50 mM Tris-HCl [pH 7.5], 150 mM NaCl, 0.1% Triton X-100) containing 1% protease inhibitor cocktail (here) and 1% Phosphatase Inhibitor Cocktail 2 (Sigma-Aldrich, P5726). After extraction, total protein was incubated with 4X Laemmli SDS Buffer (Fisher Scientific) and heated for 10 minutes at 95° C.

Proteins were separated by SDS-PAGE (10% acrylamide) and were transferred to a nitrocellulose membrane. After blocking with 1% BSA in TBS-Tween (0.01%) buffer for 1 hour at room temperature. Phosphorylation of MAP Kinases were detected by an antiphospho-p44/42 MAPK (Erk1/2) (Thr202/Tyr204) HRP-conjugated antibody (Cell Signaling Technology) and actin was detected by HRP conjugated Anti-Plant Actin Mouse Monoclonal Antiboty (3T3) (Abbkine, ABL1055). Signals were detected using SuperSignal West Pico Plus Chemiluminescent Substrate (Thermo Fisher). MAPK activation was quantified using an established ImageJ plugin (Ohgane & Yoshioka, 2019).

### Total RNA extraction for RNA-seq of Tomato Roots

Five-day-old H7996 seedlings were cut into whole root, late differentiation, and early differentiation zones using the same methods as the ROS and MPK assays. The root segments were left in water overnight to recover and then treated with 1 uM flg22, 100 nM flgII-28, or mock water. Six root samples from each segment type and treatment were pooled at 6 hpi, and the samples were ground into a powder using a mortar and pestle under liquid nitrogen. Whole root and LD samples (100 mg ± 10) or ED samples (20 mg ± 5) of root ground tissue from each sample was used for RNA extraction using Trizol (Invitrogen), following the manufacturer’s instructions. RNA purification was done with Qiagen RNeasy mini-Kit with DNase I treatment in-column treatment.

### RNA-seq

Three biological replicates (each consisting of roots from three individual plants) per genotype and treatment were subjected to Illumina RNA sequencing. Each sample averaged about 45.7 million (range from 27.1 to 66.6 million) high quality paired end reads. More than 94% of the reads were mapped to the ITAG4.1 Solanum lycopersicum reference genome, using STAR version 2.7.10.a. Gene expression was measured as the total reads for each sample that uniquely mapped to the reference gene list with summarizeOverlaps (GenomicAlignments1.34.1 and Rsamtools 2.14.0). Data was filtered for low counts such that at least three of the 12 samples had at least three counts per row. Differential gene expression analysis was performed with DESeq2 version 1.38.3. We used an FDR < 0.05 to determine differentially expressed genes. Gene ontology (GO) and KEGG analysis were performed using ShinyGo 0.80 for categories that contained less than 500 terms in their corresponding category. Heatmaps were visualized with R software version 3.4.0 package “ggplot2”.

### Statistical Analyses

Statistical analyses were conducted in R version 3.4.0. Data distribution was assessed, and tests appropriate to the distribution of the data were applied.

## RESULTS

### Whole tomato root ROS responses to PAMPs are species and cultivar-dependent and show distinct patterns among PAMPs

Leaves of different tomato cultivars recognize and respond to PAMPs with different amplitudes and durations of ROS burst (Veluchamy et al., 2014; Roberts et al., 2019;). To test whether this was true in roots, we established a whole-root ROS assay with four tomato cultivars of interest (**Table 1**). Upon treatment with 1µM flg22^Pst^, 100 nM flgII-28^Pst^ or 1µM csp22^Rsol^, cultivar H7996 and the *S. pimpinellifolium* accession LA2093 exhibited a detectable ROS burst within the first 40 minutes of treatment (**Fig. 1a,b, Fig. S1**). Yellow Pear, which lacks *SlFLS3*, and Wv700 displayed a ROS burst to only flg22^Pst^ treatment and not flgII-28^Pst^ or csp22^Rsol^. None of the cultivars responded to flg22^Rsol^ treatment, consistent with a previous report from Moneymaker tomatoes (**Fig. 1c**) (Wei et al., 2018). *S. pennellii* accession LA0716, lacking a functional *SlCORE* (Wang et al., 2016), was also used as a control for our csp22^Rsol^ peptide (**Fig. S2**).

**Fig. 1:**
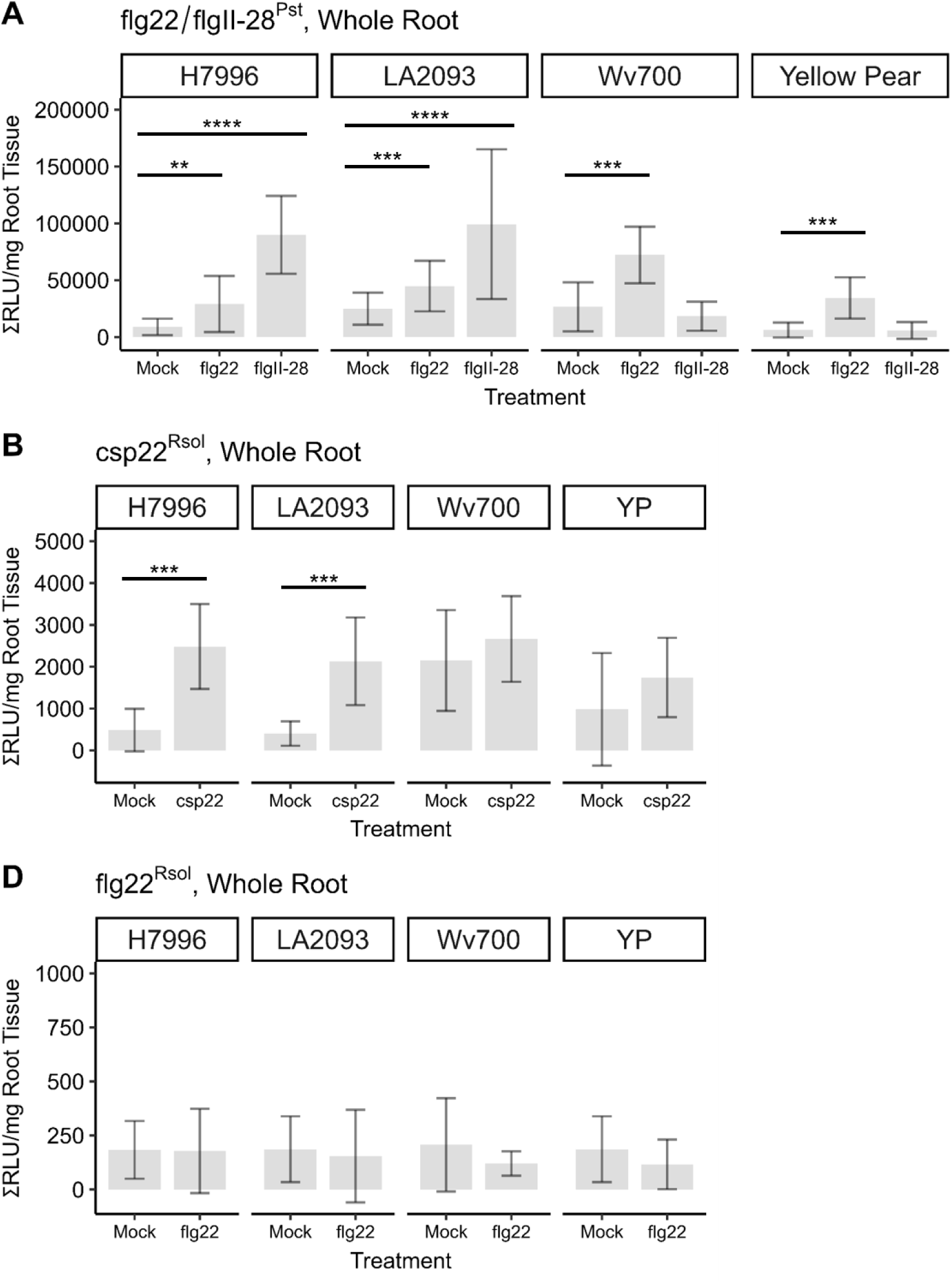
Reactive Oxygen Species (ROS) burst amplitude varies both by PAMP type and cultivar for tomato whole roots. Root samples from 5-day-old tomato seedlings of H7996, LA2093, Wv700, and Yellow Pear were treated with **(a)** 1 µM flg22^Pst^, 100 nM flgII-28^Pst^ or mock (water), **(b)** 1 µM csp22^Rsol^ or mock (water), and **(c)** 1 µM flg22^Rsol^ or mock (water). Values represent the mean ± SD from at least 18 replicates per treatment (Wilcoxon, \**p<0.05, **p<0.01, ***p<0.001, ****p<0.0001)*

H7996 and LA2093 both exhibited three distinct amplitudes of ROS burst, with the highest being flgII-28^Pst^ response, then flg22^Pst^, and lastly csp22^Rsol^ (**Fig. 1**, **Fig. S1).** The ROS burst kinetics also differed, with peak signals for csp22^Rsol^ and flg22^Pst^ at around 10 minutes post treatment and flgII-28^Pst^ elicitation at 20 minutes in H7996. flg22-induced ROS burst attenuated to basal levels by 40 minutes post treatment, while flgII-28 and csp22-induced ROS bursts were still elevated.

This data suggests varying degrees of cellular response to the bacterial PAMPs, possibly from expression levels of the respective PRR or distinct involvement of PRR-specific co-receptors and signaling components.

### Root ROS response to PAMPs is primarily located in the Early Differentiation Zone

PRR expression is highly correlated with areas of pathogen entry and colonization, including that of the stomata, stele, and sites of lateral roots. In the root, soil-borne pathogens often enter through wounded areas and natural openings. Pathogenic bacteria are shown to accumulate at the Arabidopsis elongation zone, where endodermal barriers are not yet fully established (Li et al., 2017; Tsai et al., 2023). This zone also exhibits heightened sensitivity to abiotic stresses (Dinneny et al., 2008; Iyer-Pascuzzi et al., 2011). To test whether areas of developing tissues were more sensitive to PAMPs, tomato primary roots were cut into three sections: the Meristematic Zone, the Early Differentiation (ED) Zone, and the Late Differentiation (LD) Zone. The ED Zone was characterized by the presence of visible emerging root hairs, while the LD Zone exemplified fully emerged root hairs on the primary root (**Fig. 2a**).

**Fig. 2:**
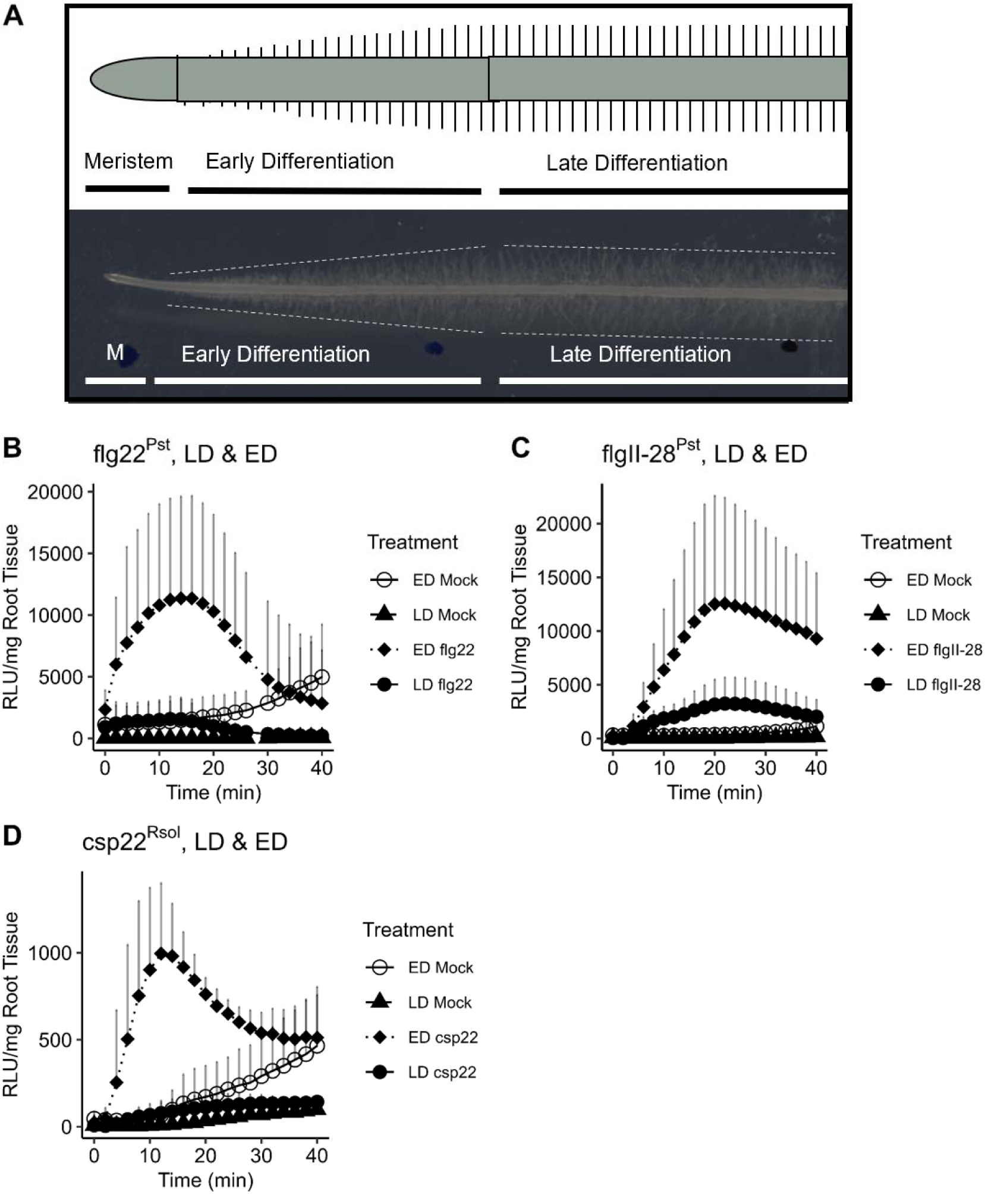
Reactive Oxygen Species Burst is primarily found in the Early Differentiation Zone. **(a)** Schematic representation of the root zones, including the Late Differentiation Zone, Early Differentiation Zone, and Meristematic/Transition Zone. H7996 treated with **(b)** 1 µM flg22^Pst^ or mock (water), **(c)** 100 nM flgII-28^Pst^ or mock (water), **(d)** 1 µM csp22^Rsol^ or mock (water). Values represent the mean ± SD from at least 6 replicates per treatment. The assay was repeated three times with similar results.

**Fig. 3.**
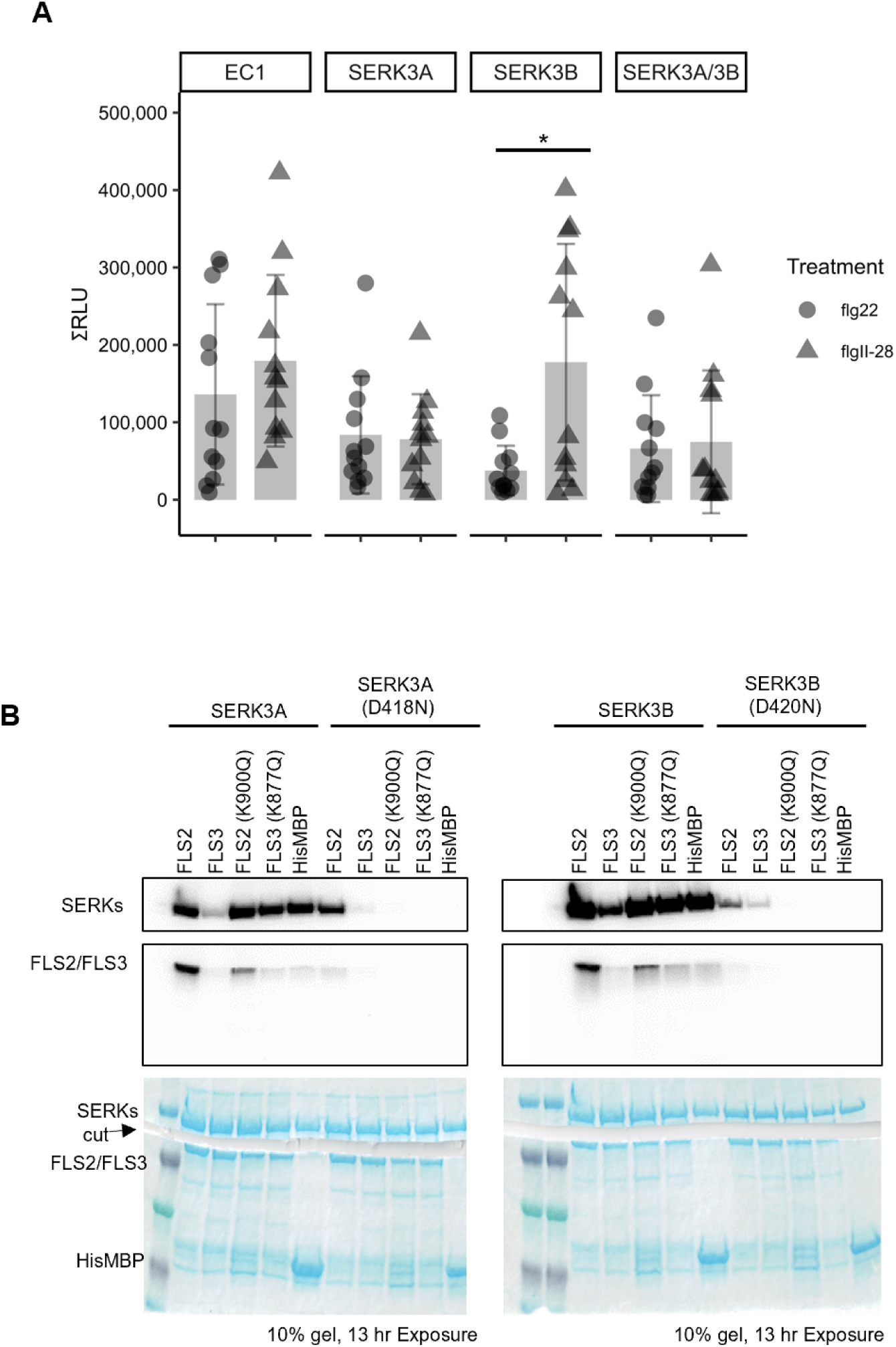
SERK3A and SERK3B interact differently with flagellin PRRs FLS2 and FLS3. **(a)** Total ROS produced through addition of peptides flg22 or flgII-28 in tomato when genes SERK3A, SERK3B, or both SERK3A and SERK3B (SERK3A/3B) are knocked down using virus-induced gene silencing (VIGS) alongside the empty control (EC1). The figure shows one representative replicate (n=4 plants of each VIGS). The experiment was repeated ten times with similar results (n=40). **(b)** *in vitro* transphosphorylation assay showing kinase activation of the cytoplasmic domains of FLS2, FLS3, SERK3A, and/or SERK3B and their kinase-inactive variants as controls (FLS2(K900Q), FLS3(K877Q), SERK3A(D418N), and SERK3B(D420N)). A protein generated from the empty vector (6x-His-MBP) was used as a negative control. FLS2 or FLS3 or their kinase-inactive variants were subjected to phosphorylation with SERK3A or its kinase inactive variant (*left panel*) or SERK3B or its kinase-inactive variant (*right panel*). Upper panels indicate ^32^P detection through a phosphor-screen. Equal protein loading is demonstrated with the Coomassie blue staining (lower panels). 10% SDS-PAGE gels were exposed to a phosphor-screen for 13 hours. The figure is from one representative replicate, and the experiment was repeated six times with similar results.

We first characterized the induction of ROS Burst for both the ED and LD Zones. In H7996, the ED zone was the primary location of ROS burst in response to flg22^Pst^ **(Fig. 2).** We then tested other varieties and found a similar burst in the other three genotypes (**Fig. S3).** Consistent with our whole root samples, both Wv700 and Yellow Pear lacked a ROS response to flgII-28^Pst^ (Fig. S3b); however, both H7996 and LA2093 exhibited ROS burst in the ED zone to flgII-28^Pst^ (**Fig. S3b**). In contrast to the lack of ROS response to csp22^Rsol^ in whole roots, both *S. lycopersicum* varieties H7996 and Yellow Pear showed a significant ROS Burst response in the ED zone compared to mock (**Fig. S3c**). Three additional *S. lycopersicum* cultivars, Brandywine, Black from Tula, and Ailsa Craig also showed significant ROS burst in the ED zone; Rutgers, however, did not (**Table S1**). Together, our data shows that the ED zone is the primary site for FLS2- and FLS3-mediated ROS burst.

### SERK3A is primarily responsible for FLS3-mediated ROS burst

To further characterize potential differences in the signaling pathway responsible for FLS2- and FLS3-mediated ROS burst, we focused on understanding the involvement of *At*BAK1 orthologs, *Sl*SERK3a and *Sl*SERK3b. Tomatoes silenced for *Sl*SERK3a, *Sl*SERK3b, or both show a severe reduction in FLS2-mediated ROS production (Peng & Kaloshian, 2014). Therefore, we asked whether *Sl*SERK3a and *Sl*SERK3b exhibited redundant functions in FLS3-mediated ROS response.

To investigate the roles of SERK3A and SERK3B in detecting flgII-28 and flg22, the expression of tomato orthologs of *SERK3A*, *SERK3B*, and both *SERK3A* and *SERK3B* (*SERK3A/SERK3B*) was knocked down in *S. lycopersicum* using virus-induced gene silencing (**Fig S4**). Knockdown in expression was confirmed using qPCR and compared to the empty control (EC1) (**Fig S4**).

Consistent with the predicted function of SERK3A and SERK3B as the presumed orthologs of Arabidopsis BAK1 (AtBAK1), knocking down *SERK3A, SERK3B,* and *SERK3A/SERK3B* reduced the flg22^Pst^ ROS burst compared to EC1 (**Fig. 3a**). However, while there was a reduced flgII-28^Pst^ ROS burst for the *SERK3A* knockdown, the flgII-28^Pst^ ROS burst in *SERK3B* showed no difference compared to the empty control. As expected, the *SERK3A/SERK3B* double knockdown also showed a reduced ROS burst for flgII-28^Pst^. This suggests that SERK3A is necessary and sufficient for immunity activation by FLS3, and the SERK3B differentially interacts with FLS2 vs FLS3 in tomato.

### FLS2 and FLS3 interact differently with SERK3A and SERK3B *in vitro*

It is possible that the differences in ROS burst could be attributed to differences in phosphorylation of SERK3A and SERK3B by their PRR co-receptors FLS2 and FLS3. It was previously reported that tomato FLS2 and FLS3 have stronger kinase activity than AtFLS2, with FLS3 having stronger kinase activity compared to FLS2. Only FLS3 could transphosphorylate a generic substrate, myelin basic protein (Roberts et al., 2020). These differences, along with the VIGS ROS burst data, suggest that FLS2 and FLS3 interact differently with the SERKs.

To test whether there are transphosphorylation differences between the PRRs and the SERK co-receptors, we made recombinant proteins in BL21 *Escherichia coli* that contained the cytoplasmic domains of FLS2 or FLS3, which are required for kinase activity (Roberts et al., 2020), and the cytoplasmic domains of SERK3A and SERK3B, expressed in the pDEST-HisMBP vector (containing a N-terminal 6xHis-MBP tag). For negative controls, we generated mutants with substitutions in their ATP binding domains, FLS2(K900Q), FLS3(K877Q), SERK3A(D418N), and SERK3B(D420N), and included a vector that expressed a short unstructured *E. coli* sequenced to permit expression of the His-MBP protein in the pDEST-HisMBP vector. In the *in vitro* kinase assays, we added either FLS2 or FLS3 and SERK3A or SERK3B, and their associated kinase inactive variants (**Fig. 3b**). When FLS2 and SERK3A are placed in the same reaction, both FLS2 and SERK3A are phosphorylated. However, when FLS3 and SERK3A are placed in the same *in vitro* kinase reaction together, FLS3 (which typically has stronger kinase activity than FLS2) and SERK3A both show a severe reduction in phosphorylation. When the kinase-inactive FLS2 (FLS2(K900Q)) and SERK3A are added together, FLS2(K900Q) is phosphorylated but SERK3A phosphorylation is reduced compared to the FLS2 wildtype reaction. With the kinase-inactive FLS3 (FLS3(K877Q)), FLS3(K877Q) is strongly phosphorylated and SERK3A has weak phosphorylation. Conversely, when the kinase active versions of FLS2 and FLS3 are added with the kinase-inactive SERK3A(D418N), FLS2 phosphorylation is maintained but SERK3A(D418N) phosphorylation is severely reduced compared to the wildtype SERK3A. Similar to the wildtype, FLS3 and SERK3A(D418N) phosphorylation are both very weak when added together.

A similar pattern is observed when the PRRs or their variants are added with SERK3B or the kinase inactive version; FLS2 and SERK3B are both phosphorylated, FLS3 and SERK3B are very weakly phosphorylated, FLS2(K900Q) and SERK3B are phosphorylated, and FLS3(K877Q) is phosphorylated but SERK3B is not. FLS2 is weakly phosphorylated when added with SER3B(D420N), whereas FLS3 is not, and SERK3B(D420N) is weakly phosphorylated.

Together, these data suggest that FLS2 and FLS3 interact differently with SERK3A and SERK3B.

### PTI driven MPK activation is PRR-specific and is primarily located in the Early Differentiation Zone

The Arabidopsis MAPK3/MAPK6 homologs, SlMPK1/2/3, are signaling proteins in the tomato immune pathway upstream of defense gene transcriptional regulation (Pedley et al., 2004; Stulemeijer et al., 2007; Wilmann et al., 2014). To test whether these signaling proteins were conserved downstream of FLS3 and CORE, we first observed MPK1/2/3 phosphorylation of eight-week-old leaf tissue in H7996 upon treatment with 1µM flg22^Pst^, 100 nM flgII-28^Pst^ or 1µM csp22^Rsol^. As expected, flg22^Pst^ treatment resulted in activation of both MPK1/2 (45 kDa)

and MPK3 (42 kDa) (**Fig. 4a**) (Pedley et al., 2014). In contrast, flgII-28^Pst^ exhibited MPK phosphorylation for MPK1/2, but not MPK3, and treatment with csp22^Rsol^ did not result in any phosphorylation. These data suggest that immune signaling pathways downstream of PRRs diverge in tomato.

**Fig. 4.**
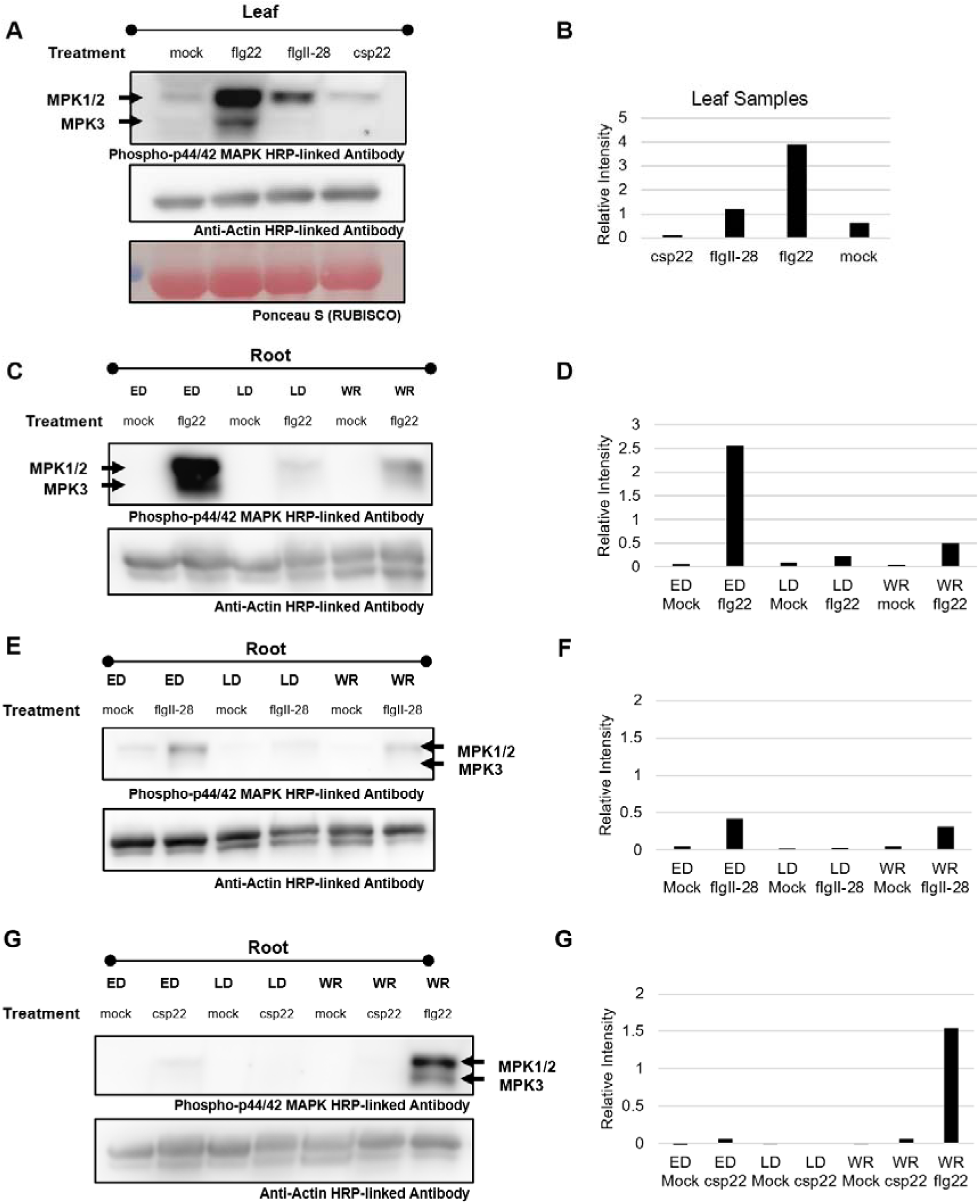
MAPK phosphorylation in tomato leaf and root tissues upon treatment with various PAMPs. **(a**) Eight-week-old leaf samples treated with mock (water), 1 µM flg22, 100 nM flgII-28, or csp22. **(b)** Quantification of MAPK phosphorylation in leaf samples from 4a, normalized to actin. **(c)** Root sections representing LD, ED, and WR treated with mock (water) or 1 µM flg22. **(d)** Quantification of MAPK phosphorylation in root sections from 4c, normalized to actin. **(e)** Root sections treated with mock (water) or 100 nM flgII-28. **(f)** Quantification of MAPK phosphorylation in root sections from 4e, normalized to actin. **(g)** Root sections treated with mock (water) or csp22. **(h)** Quantification of MAPK phosphorylation in root sections from 4g, normalized to actin. Phosphorylation was assessed by western blot using Phospho-ERK1/2 HRP-linked antibody (CellSignaling, #8544). Total proteins were detected by Anti-Actin HRP- linked Antibody (Abbkine). A Bradford assay was also used for equal protein loading. The assay was repeated three times with similar results.

We next asked whether MPK1/2/3 phosphorylation occurred in the root, and if so, whether it followed our ROS burst data and primarily occurred in the ED zone. Thus, we observed MPK phosphorylation of the ED, LD and whole root after treatment with 1µM flg22^Pst^, 100 nM flgII- 28^Pst^ or 1µM csp22^Rsol^ in five-day old tomato seedlings. Consistent with our ROS data, the ED Zone showed heightened MPK phosphorylation when compared to the LD Zone or whole root for both flg22^Pst^ and flgII-28^Pst^ treatment (**Fig. 4c-f**). Similarly to the ED ROS data for csp22^Rsol^, the strength of the PTI response was far lower, if not absent, in tomato ED, LD, and WR sections (**Fig. 4g,h**). In parallel with the developmental-specificity of PTI-driven ROS burst, these data not only suggest that the ED zone is the primary location for PTI initiation and response, but also that ROS burst and MPK phosphorylation – representative of two distinct downstream pathways – are differentially controlled within the receptor complex.

### Transcriptional reprogramming after PAMP treatment is heightened in the ED Zone

To further understand the link between developmental specificity of PTI initiation and subsequent transcriptome modifications, we used root sections of the PAMP-responsive cultivar H7996 treated with 1µM flg22^Pst^ or 100 nM flgII-28^Pst^. Whole root, ED, and LD sections were cut, washed, and left overnight before PAMP or water treatment (Modified from Wei et al., 2018). At 6 hours post treatment, roots were collected for RNA extraction, sequencing, and subsequent analysis using DESeq2 for identification of differentially expressed genes (DEGs). Whole root samples treated with flg22^Pst^ or flgII-28^Pst^ differed in the number DEGs than either of the treated ED samples. In the flg22^Pst^ whole root samples, 3836 and 2145 genes were up- regulated and downregulated, respectively, while the ED samples exhibited 2959 upregulated and 3835 downregulated genes (**Fig. 5a,b)**. In the flgII-28^Pst^ whole root samples, 248 genes were upregulated while 221 genes were downregulated in comparison to the ED Zone’s 1843 upregulated and 1910 downregulated genes (**Fig. 5a,b).** Only 144 upregulated and 132 downregulated genes were shared between whole root treatments, while 1496 upregulated and 1513 downregulated genes were shared between treatments for ED samples. The majority of the DEGs found in the whole root samples for each treatment were not identified in our ED samples (**Fig. 5c)**. The identification of genes distinctly upregulated in the ED shows that transcriptional regulation in the whole root is not reflective of the ED response and is consistent with our data showing the ED exhibits a distinct PTI response. In addition, the increased number of PTI-associated DEGs for flg22^Pst^ compared to flgII-28^Pst^ is consistent with our findings that, overall, flg22^Pst^ treatment results in more prominent transcriptional reprogramming.

**Fig. 5.**
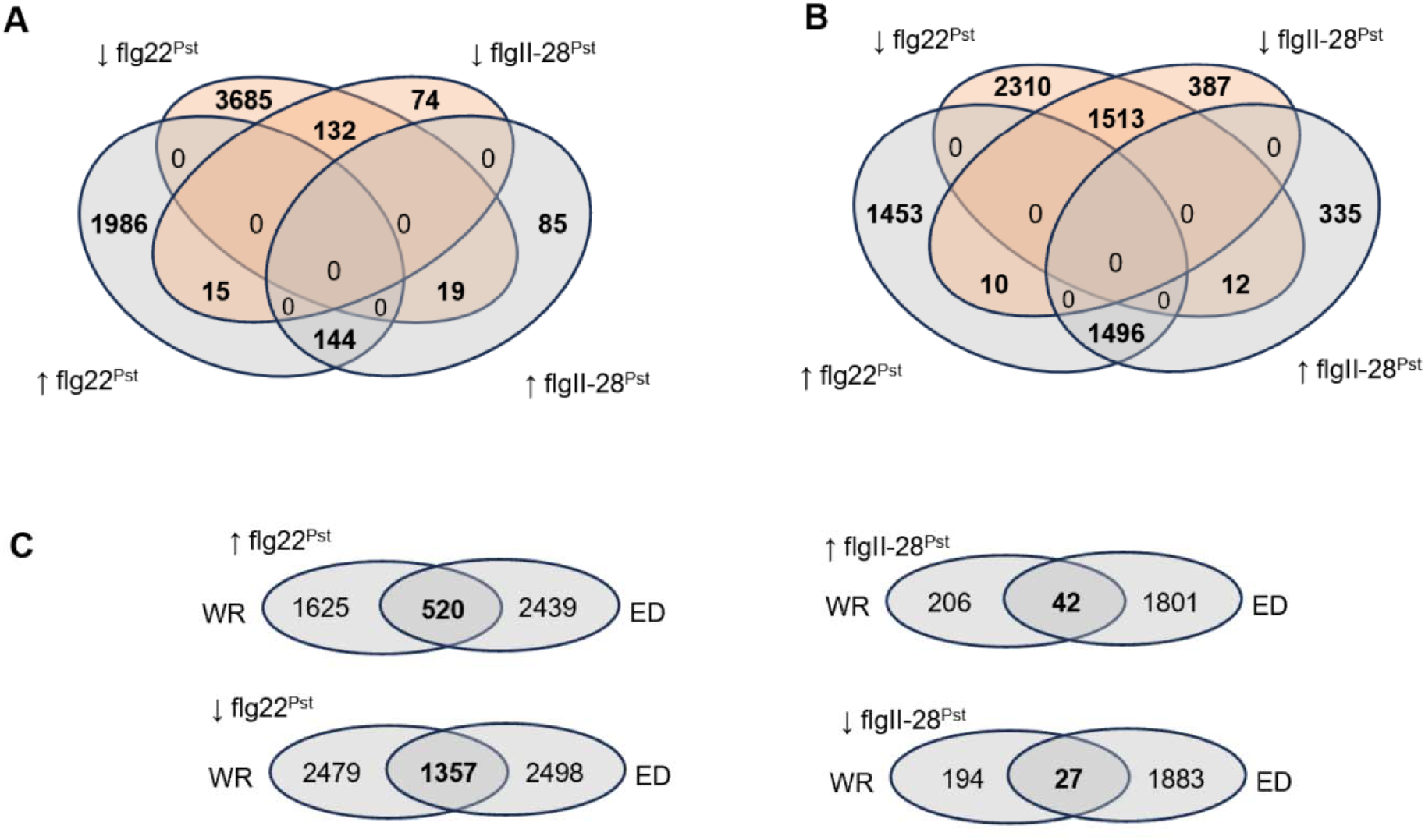
Differentially expressed genes six hours after treatment with 1 uM flg22 or 100 nM flgII-28. Venn diagram depicting both up- and downregulated DEGs for **(a)** whole root or **(b)** Early Differentiation zone samples after treatment with flg22 or flgII-28. **(c)** Overlap in DEGs for Whole Root and Early Differentiation Zone samples. DESeq2, p-adj < 0.05.

To more accurately understand the function of the DEGs found in our analysis, we performed a KEGG Pathway analysis and GO Biological Function analysis with the ShinyGO toolkit 0.80. Our GO Biological Function analysis found that transcription for genes involved in plant-pathogen immune responses was increased in the ED Zone. Of the top 20 KEGG categories (False Discovery Rate < 0.05) for each treatment, 13 categories were shared between flg22 and flgII-28 in the ED Zone, including “Cellular Response to Chemical Stimulus” “Intracellular Signal Transduction,” “Response to Biotic Stimulus,” “Response to Other Organism,” “Response to External Biotic Stimulus,” “Biological Processes Involved in the Interspecies Interaction Between Organisms,” (Supplemental Table 1). Between treatments, the ED zone showed an increased number of genes for flg22 in each of the categories compared to flgII-28 treatment **(Fig. S5a,b)**. FlgII-28 response initiated the exclusive transcription of two Ethylene-Responsive Transcription Factors (ERFs) *Solyc05g051200* and *Solyc09g066350* as compared to two distinctly upregulated ERFs (*Solyc04g012050* and *Solyc06g068830*) and a number of ethylene receptors upon flg22 perception. Consistent with PTI response in other species, transcripts associated with cell wall and cytoskeleton organization were downregulated in response to both PAMPs (“Cell Wall Organization or Biosynthesis”) (Wang et al., 2022).

In comparing the GO Biological Function categories most upregulated between ED and whole root samples, we found that flg22 treatment shared six of the top twenty categories of upregulated genes (“Cellular Response to Chemical Stimulus,” “Intracellular Signal Transduction,” “Cellular Response to Organic Substance,” “Response to Oxygen Containing Compound,” and “Cellular Response to Hormone Stimulus”) **(Fig. S5a,c)**. These tended to be broad, less specific categories, and in each category, the shared number of genes between whole root and ED samples were low. For example, the category “Cellular Response to Chemical Stimulus” exhibited 40 DEGs for whole root samples and 72 for ED samples; however, the two root types only shared 13 genes. The same was true for flgII-28 whole roots, with an even lower number of genes upregulated per category and no overlap between ED and whole root samples **(Fig. S5b,d)**. Together, this functional analysis further supports a more complex flg22 response compared to flgII-28 and provides evidence that the whole root is not representative of the ED zone’s sensitivity to PTI.

We next asked whether responses in tomato roots were similar to commonly-associated PTI genes that included PRRs, co-receptors, and downstream signaling elements (Gomez-Gomez et al., 2000; Li et al., 2015; Wei et al., 2018; Hind et al., 2016; Zhang et al., 2020). Out of the twelve PTI-associated genes, only one, *SlRbohB*, showed significant upregulation in the whole root for flgII-28^Pto^ treatment; five of the PTI-associated genes were significant for flg22^Pst^- treated whole roots **(Fig. 6)**. In contrast, eleven of twelve PTI-associated genes were differentially expressed after flg22^Pst^ treatment in the ED samples, and seven of the twelve transcripts were differentially expressed after treatment with flgII-28^Pst^ (**Fig. 6**). Interestingly, five genes (*FLS2.1*, *MPK3*, and *RbohB, and WRKY33A/B*) were significantly downregulated in whole root samples while they were significantly upregulated in ED samples.

**Fig. 6.**
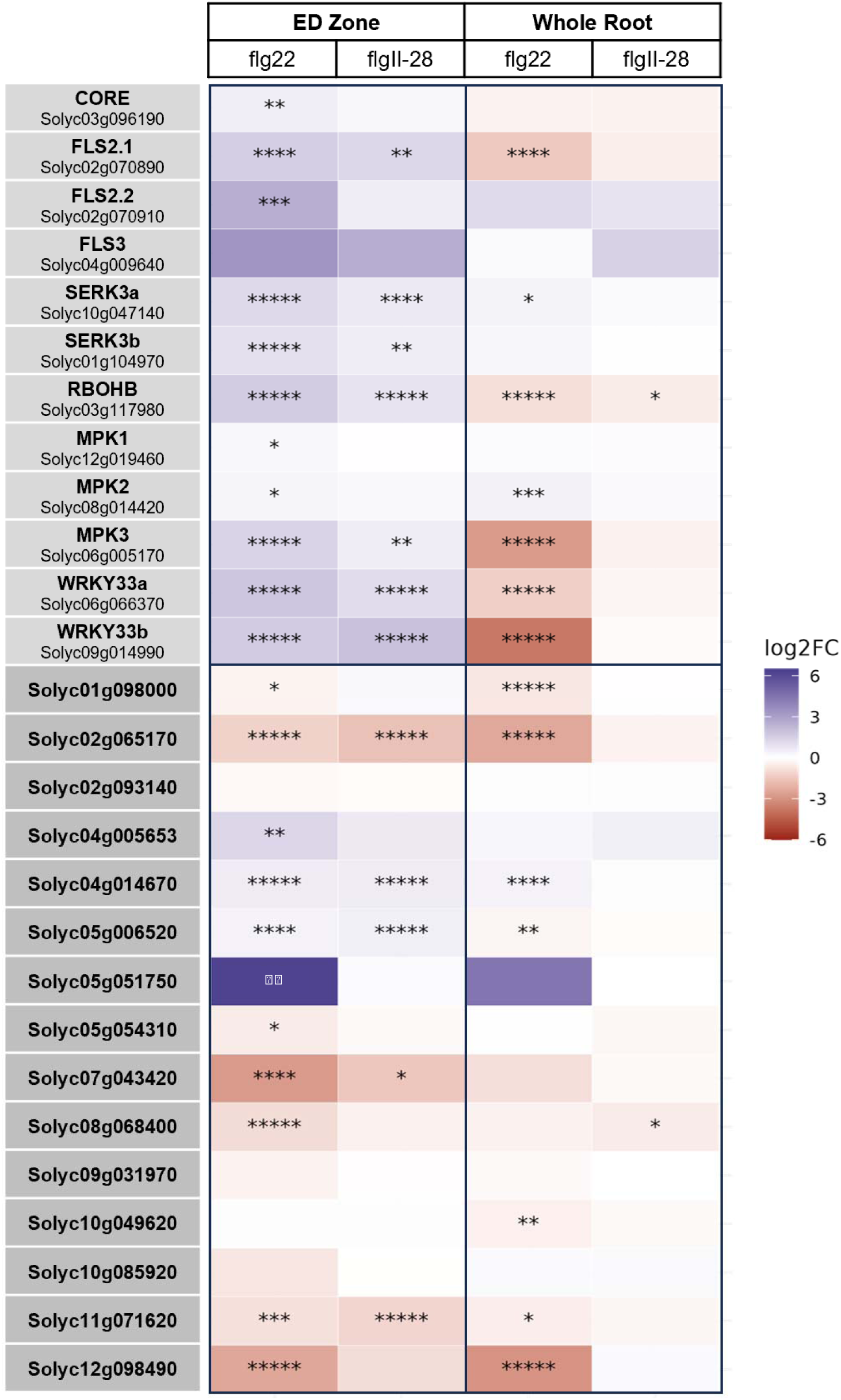
Expression of genes from the RNAseq dataset that encode for proteins directly associated with the PTI signaling pathway as well as PTI-marker gene candidates from Yu. et al (2021). The colors of the graph represent the Log2FC, while significance is shown through the p–adj values: < 0.05*, 0,01**, 0.001***, 0.0001****, 0.00001***** .

We next evaluated fifteen potential PTI marker genes identified in the proteomic analysis by Yu et al. (2021), which had not been detected in previous, whole-leaf and whole-root transcriptional studies. Upon comparison of the gene expression levels within our whole-root data, a single gene (*Solyc08g068400*) was significantly repressed in the whole root samples treated with flgII-28^Pst^ and seven were significant for whole root samples treated with flg22^Pst^ (**Fig. 6**). In contrast, ED zones treated with flgII-28^Pst^ showed significant differential expression for 5 of the 15 candidate PTI marker genes and treatment with flg22^Pst^ resulted in significant differential expression for 11 of the 15 candidates in the ED zone. All 5 DEGs from flgII-28^Pst^ treatment were found within the flg22^Pst^ DEGs. Together, these results support our understanding of PTI specificity in the ED zone and identify five candidate PTI marker genes (*Solyc02g065170, Solyc04g014670, Solyc05g006520, Solyc07g043420, Solyc11g071620*) for both proteomic and transcriptomic studies.

### Early root growth inhibition is a result of flg22-mediated PTI, but not of flgII-28 or csp22

Our results revealed differences between early tomato root responses to flg22 and flgII-28 and signaling downstream of FLS2 and FLS3. To test whether the differences in immune signaling resulted in different phenotypic outcomes, we tested the impact of each peptide on root growth. Although prolonged flg22 exposure leads to seedling growth inhibition (Gomez-Gomez, 2000), we hypothesized that the robust transcriptional response in the tomato root ED may have an observable phenotypic outcome after transient exposure to flg22. Given the relative differences in transcriptional reprogramming between flg22^Pst^ and flgII-28^Pst^ treatment, we reasoned that tomato roots would show a more prominent phenotypic response to flg22^Pst^ treatment than flgII- 28^Pst^. We hypothesized that csp22^Rsol^ treatment would have a minor influence on root growth, similar to that of ROS burst amplitude and attenuation.

Upon a single treatment with 1µM flg22^Pst^ root growth in each of the four cultivars tested was temporarily inhibited for the first 24 hours post inoculation (hpi) but recovered to that of mock by 48 hpi (**Fig. 7a**). Treatments of both 100 nM flgII-28^Pst^ and 1µM csp22^Rsol^ on all four cultivars failed to elicit temporary growth inhibition at both 24 and 48 hpi (**Fig. 7b,c).** As a control, roots were treated with flg22^Rsol^ (**Fig. 7d**). The absence of temporary root growth inhibition for flgII-28^Pst^ and csp22^Rsol^ strengthens our hypothesis that FLS2-mediated PTI is more sensitive than other bacterial-peptide recognition by respective PRRs and that downstream elements of PTI are independent yet overlapping.

**Fig. 7.**
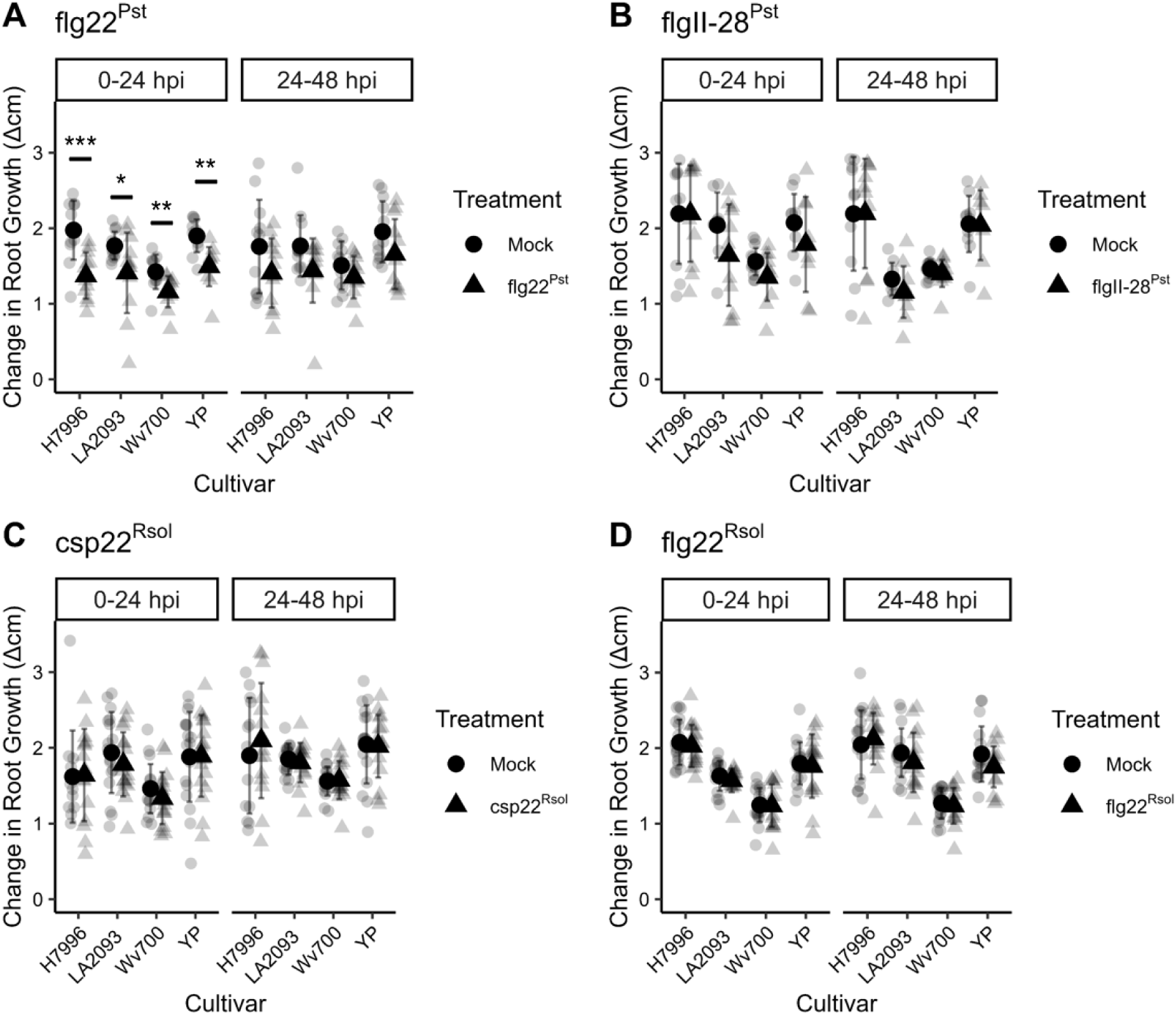
Temporary root growth inhibition is observed for flg22^Pst^ treatment, but not flgII- 28^Pst^, csp22^Rsol^, or flg22^Rsol^. Change in root growth (cm/24 hour) for tomato roots of cultivars H7996, LA2093, Wv700, and Yellow Pear from 0-24 hours and 24-48 hours. Tomato seedlings treated with **(a)** 1 µM flg22^Pst^ or mock (water), **(b)** 100 nM flgII-28^Pst^ or mock (water), **(c)** 1 µM csp22^Rsol^ or mock (water), and **(d)** 1 µM flg22^Rsol^ or mock (water). Values represent the mean ±SD from at least 12 roots per treatment (Wilcoxon, \**p<0.05, **p<0.01, ***p<0.001, ****p<0.0001)*

To test whether this same temporary growth inhibition and recovery occurred in other FLS2- mediated PTI events, we performed a root growth assessment on Arabidopsis (Col-0) seedlings with the same single PAMP flood treatment. Notably, a temporary FLS2-mediated root growth inhibition for flooded Arabidopsis seedlings did not occur until 48 hours post treatment (**Fig. S4a**). Similar to tomato, the Arabidopsis seedlings resumed normal growth rates just 24 hours later. Overall, our experiments indicate that the strength of root growth inhibition to a single PAMP treatment varies among elicitors, and tomato root response and recovery to a single flg22^Pst^ elicitation occurs more rapidly than that of Arabidopsis.

### Temporary root growth inhibition (RGI) is independent of ROS burst in tomato root PTI

In Arabidopsis, PAMP-induced prolonged RGI is independent of the NADPH oxidase RBOHD (Lu et al., 2009; Shinya et al., 2014; Tran et al., 2020). In tomato, the NADPH oxidase *Sl*RbohB has been linked to PTI-derived ROS burst, but it remains unclear whether *Sl*RbohB and RGI are directly linked. We found that flgII-28^Pst^ induced a ROS burst with higher amplitude and longer attenuation compared to roots treated with flg22^Pst^. However, roots treated with flg22^Pst^, but not flgII-28^Pst^, exhibited a temporary RGI. These results prompted us to ask whether the PTI-derived ROS burst and temporary RGI are independent processes in tomato.

To examine whether ROS production and temporary RGI are independent, we performed RGI assays using the NADPH oxidase inhibitor diphenyleneiodonium chloride (DPI) alongside flg22^Pst^ treatment for tomato cultivars H7996 and LA2093. We first identified the minimum concentration of DPI needed to fully inhibit the ROS burst response (**Fig. S6)**. Using this concentration (1 µM), we pre-treated tomato seedlings with either DPI or a mock solution before applying 1µM flg22^Pst^. Despite the DPI treatment, temporary root growth inhibition was still observed in both H7996 and LA2093 (**Fig. 8A-B),** suggesting that temporary RGI was not dependent on NADPH-produced ROS.

**Fig. 8.**
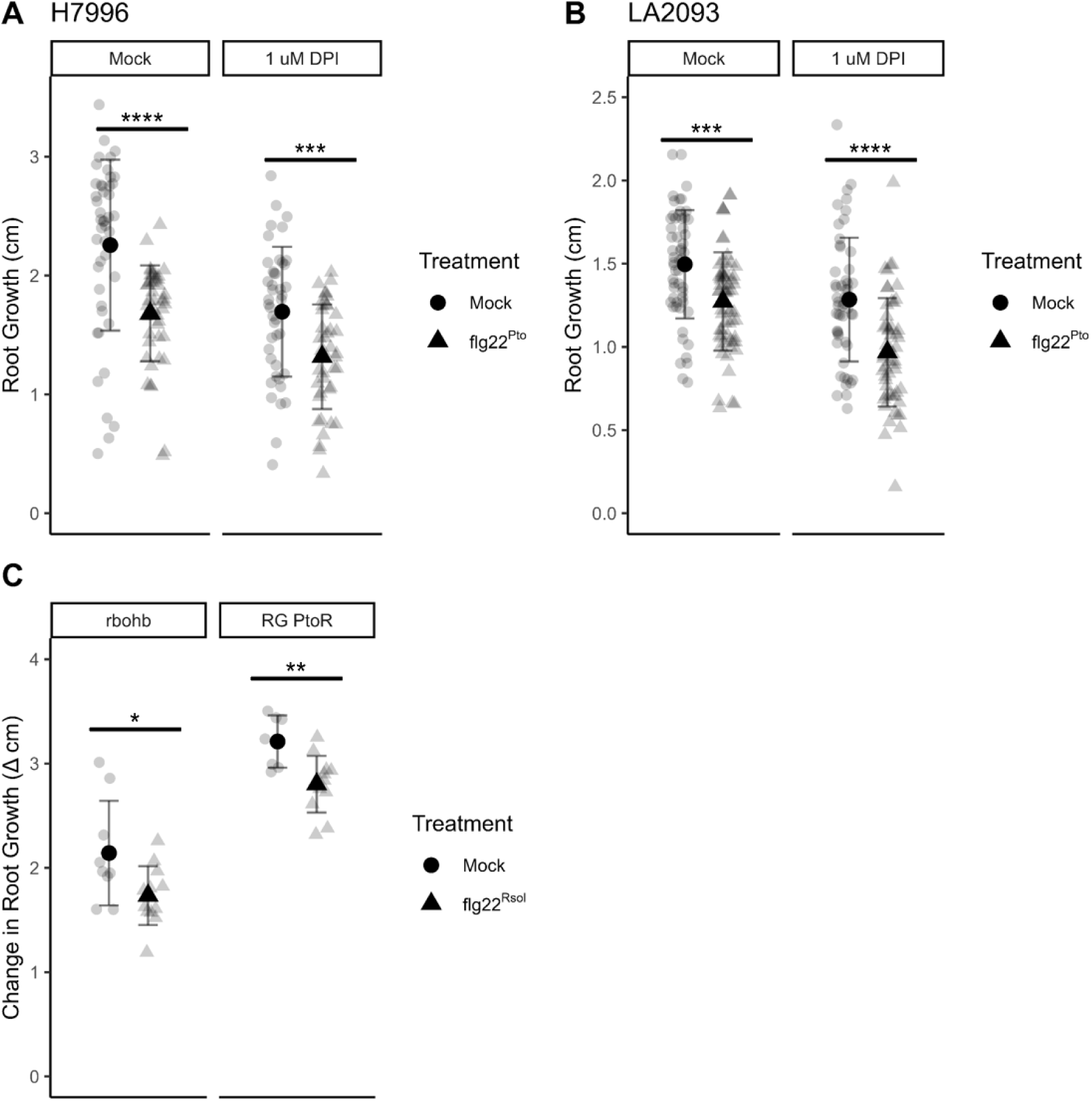
Temporary RGI is independent of ROS burst in tomato root PTI response. Change in root growth (cm/24 hour) for tomato from 0-24 hours and 24-48 hours. Five-day-old tomato seedlings of (**A**) H7996 and (**B**) LA2093 were treated with 1 µM DPI or mock (water) four hours prior to 1 µM flg22^Rsol^ or mock (water) treatment. Values represent the mean ±SD from at least 36 replicates per treatment. (Student’s t-test, *p<0.05, **p<0.01, ***p<0.001, ****p<0.0001)

To further investigate the independence of ROS burst and temporary RGI, we repeated the initial single-treatment growth inhibition experiment on seedlings with a point mutation in *Sl*RbohB, leading to a frameshift in exon 1. The *Slrbohb* line displayed an abolishment in ROS response upon treatment with 100 nM flg22^Pae^ (**Fig. S8)**. Upon treatment with 1 uM flg22^Pst^, the *Slrbohb* lines exhibited temporary RGI at 24 hours compared to mock treatment (**Fig. 8C).** Together, these results strongly suggest that temporary RGI and ROS burst function independently in *Sl*FLS2-mediated PTI.

We next asked whether FLS2-mediated temporary RGI in Arabidopsis was dependent on ROS. We used both *AtrbohD* and *AtrbohD*/*rbohF* knockout lines to measure RGI in response to 1µM flg22 treatment. The wild-type (Col-0) and the *rbohD* mutant lines exhibited RGI between 24 and 48 hpi, similar to previous experiments with flg22^Pst^ (**Fig. S4B)**. These findings are consistent with the independence of ROS burst and FLS2-mediated temporary RGI in Arabidopsis (Lu et al., 2009; Shinya et al., 2014; Tran et al., 2020).

## DISCUSSION

Pattern-triggered immunity (PTI) plays a crucial role in the innate immune response of plants, including tomato, where it is activated by the recognition of conserved microbial patterns through pattern recognition receptors (PRRs). These receptors, integral to the detection of and defense of pathogens, have been successfully transferred within and among species, showing promise in broad-spectrum resistance strategies for crop protection. However, to effectively engineer crops for broad-spectrum resistance, we must first understand how each PRR functions in its plant of origin. Our work aims to understand the PTI dynamics of three PRRs in tomato roots responsible for the recognition of three distinct bacterial-derived PAMPs. We first set out to compare hallmark elements of PTI between these PRRs, finding that both responses to flgII- 28 and csp22 were distinct from the flg22 response in the short-term MPK1/2/3 activation and temporary root growth inhibition. For the flagellin-derived PAMPs, we also noted differences in flgII-28 recognition that included a primary reliance on SERK3a, but not SERK3b, for the initiation of ROS burst and an overall lower level of transcriptional reprogramming. Further, we found that signature elements of PTI, such as MPK phosphorylation and ROS burst, are development-specific in the root, occurring primarily in the Early Differentiation Zone.

Consistent with this, we found that the ED Zone is also a hotspot for defense gene activation upon PTI activation.

Together, our data shows that individual LRR-RLKs initiate distinct PTI responses in tomato roots that are highest in the root’s developing tissues.

### FLS2, FLS3, and CORE response exhibit natural variation in tomato roots

Variation in PTI response among cultivars of *S. lycopersicum* has been well documented within leaf samples, including that of differences in amplitude and attenuation of ROS formation (Veluchamy et al., 2014; Roberts et al., 2019; Moroz & Tanaka, 2020). Our study reveals key details of PTI signaling in below ground tissues, including that the ROS production also varies in amplitude and attenuation within tomato roots. Specifically, flgII-28^Pst^ induces a stronger response compared to flg22^Pst^, despite the lower concentration of flgII-28^Pst^ (100nM) to flg22^Pst^ (1 μM). The flgII-28^Pst^ response showed a more prolonged attenuation than flg22^Pst^, with ROS production returning to basal levels after over 60 minutes, whereas ROS levels in flg22^Pst^-treated samples returned to baseline after 40 minutes. On the contrary, treatment with 1uM csp22^Rsol^ elicited only 1/20^th^ of the ROS burst in H7996 as flg22. Considering the amplitude of ROS response for flgII-28^Pst^, we hypothesized that temporary RGI for the four cultivars would be stronger after flgII-28^Pst^ treatment than after flg22^Pst^. We were surprised to see that flgII-28 samples, in fact, showed no temporary RGI in any of the four cultivars. The absence of temporary and RGI for flgII-28^Pst^ treatment despite the stronger ROS response suggests that just with *At*FLS2, flgII-28 recognition drives separable PTI responses (Colianni et al., 2021).

For csp22 response, our whole root H7996 and LA2093 samples exhibited a strong ROS burst, while Wv700 and Yellow Pear lacked a significant ROS. This was different than our results for ED sections, where H7996 and Yellow Pear showed significant ROS burst. The contrasting csp22 responses in LA2093 between whole root and ED zone suggest that CORE may not have ED-specific involvement in PTI response. Due to the speculation of both age-specific (Wang et al., 2016; Dodds et al., 2023) and development-specific *Sl*CORE expression, we further investigated whether the heightened csp22^Rsol^ ROS response was seen in four additional varieties of tomato (*S. lycopersicum*) and confirmed that, just as with flg22 and flgII-28 treatment, CORE- mediated ROS response was present in the ED Zone for three of the four samples. If not solely age dependent, the lack of ROS response to csp22 made us question the role of CORE-mediated PTI in quantitative resistance strategies.

These data show that tomato cultivars not only show peptide-specific ROS burst amplitude and attenuation, but also exhibit natural genetic variation in root ROS responses similar to leaf tissues. Additionally, the strength of ROS and RGI responses are not directly correlated. The separation in response type and strength alludes to the existence of tightly regulated, PRR specific downstream pathways.

**Tomato roots exhibit distinct, but overlapping, PTI responses to immunogenic peptides.** The Pattern Triggered Immunity (PTI) model was originally established in Arabidopsis and has sense served as a foundation in understanding plant immune responses, particularly in the role of LRR-RLK FLS2 and its associated counterparts: co-receptor *At*BAK1 (Li et al, 2002; Sun et al, 2013), receptor-like cytoplasmic kinase *At*BIK1 (Lu et al., 2010; Zhang et al., 2010), mitogen-activated protein kinases *At*MAPK3/6 (Asai et al., 2002) and NADPH oxidase *At*RBOHD (Li et al., 2014) and WRKY 33 (Zipfel et al., 2004). While these findings on FLS2 activation have helped to drive PTI-based bioengineering strategies for broad spectrum resistance, PRRs such as tomato FLS3 and CORE are not found within the Arabidopsis genome (Ngou et al., 2022). FLS3 is only found within Solanaceous plants (Clarke et al., 2013) and CORE is found in both Solanaceous plants including *N. benthamiana* (Wang et al., 2016).

Upon our initial discovery that FLS2 and FLS2-mediated PTI result in characteristic differences between ROS burst and MPK activation compared to FLS3, we hypothesized that these downstream elements of PTI must be differentially regulated by complex co-receptors or RLCKs in the PTI pathway. *At*BAK1 homologs *Nb*BAK1, *Sl*SERK3A, and *Sl*SERK3B are known to form a complex with FLS2 in *N. benthamiana* and tomato (Peng & Kaloshian, 2014; Hind et al., 2016). In accordance with previous findings, our results show an increased amplitude and prolonged attenuation of flgII-28 ROS response compared to that of flg22 (Roberts et al., 2020) which led us to believe that the activation of SERK3a and SERK3b may differ for FLS3 response. Our results here show that silencing of SERK3a, but not SERK3b, resulted in a significant decrease of ROS response by tomato FLS3 (Fig. 8).

Independent of ROS response, our results show that PTI initiates a downstream activation of MPKs leading to transcriptional reprogramming and the upregulation of defense genes in tomato roots. Like that of Arabidopsis, flg22 perception activates *At*MAPK3 and *At*MAPK6 homologs *Sl*MPK3 and *Sl*MPK1/2, respectively (Fig. 3) (Pedley, 2004; Willman, 2014). FlgII-28 perception in tomato roots primarily activates SlMPK1/2 phosphorylation (Fig. 3), which is similar to FLS3 perception in *Solanum tuberosum* (Moroz & Tanaka, 2020). This differs from the activation of MAPK1/2/3 seen in tomato protoplasts (Hind et al., 2016). Interestingly, *S. tuberosum* exhibits MPK phosphorylation upon flgII-28 perception, but not flg22 (Moroz & Tanaka, 2020). Although, it is possible that the MPK3 is present in roots but too low to be detected. Unlike flg22 and flgII-28, csp22 treatment resulted in a lack of MPK1/2/3 phosphorylation for five-day-old tomato roots or eight-week-old tomato leaves, suggesting that though *Sl*CORE transiently expressed in *N. benthamiana* leaves phosphorylate *Nb*MAPK3 and *Nb*MAPK6, significant MPK phosphorylation upon csp22 recognition is absent in tomato roots (Wei et al., 2016). Together, these data suggest that while there are conserved elements in PTI signaling across species, there are also unique aspects within the Solanaceae family that have evolved to optimize pathogen recognition and defense mechanisms.

In Arabidopsis, the RLCK *At*BIK1 is often referred to as a central regulator underlying PTI, integrating signals from multiple PRRs and responsible for the activation of NADPH oxidase *At*RBOHD (Li et al., 2014; Bi et al., 2018). In tomato, functional divergence has resulted in no direct *At*BIK1 homolog; however, a handful of RLCKs in the tomato genome have known functions and interact with FLS2 and FLS3. RLCKs involved in flagellin-derived PTI response are not well characterized beyond that of *Sl*TPK1b (FLS2) and *Sl*FIR1 (FLS2/FLS3) (AbuQamar et al., 2008; Sobol et al., 2023). Mutations in *Sl*FIR1 exhibit lower levels of ROS upon treatment with both flg22 and flgII-28; in contrast, levels of MPK phosphorylation in tomato leaves were unaffected for both treatments. Whether this change in downstream elements is driven by the direct phosphorylation of SERKs by FLS2/ FLS3 or by differential phosphorylation of downstream RLCKs is not fully understood. Characterization of additional RLCK homologs in tomato will provide a clearer picture of PTI regulation in tomato.

### Tomato root PTI is specific to the early differentiation zone

PTI is a tightly regulated process, with PRRs exhibiting both spatial and developmental specificity. For example, the *EF-Tu* receptor in Arabidopsis is expressed only in above-ground tissues (Wyrsch et al., 2015), while At*FLS2* demonstrates tissue specific expression, particularly in the stele, stomata, and lateral roots (Beck et al., 2014). This expression of PRRs is highly correlated with entry sites of potential pathogens such as natural openings or wounds within the plant tissues. For soilborne pathogens such as *Ralstonia solanacearum*, these natural openings are thought to include developing root tissues and lateral root emergence sites (McGarvey et al., 1999; Caldwell et al., 2017). Our data showed that the molecular components of PTI are most prevalent in the early differentiation (ED) zone, a region in which root cells are elongating and initiating the process of differentiation.

In tomato, the *At*RBOHD homolog *Sl*RbohB is not only responsible for PTI-driven ROS burst, but also contributes to the regulation of primary root elongation and development (Zhou et al., 2020). Our results showed that the ED zone – marked by the absence of fully developed root hairs – is the primary location of ROS response for flg22^Pst^, flgII-28^Pst^, and csp22^Rsol^. The ED appeared to be a ‘hotspot’ for PTI responses, as ROS activity was not the only development-specific PTI response. The ED zone showed significant upregulation in MPK activation compared to the whole root, and the LD zone lacked a MAPK response. Together, the presence of both ROS and MPK activation in the ED tissues are consistent with the localization of PTI sensitivity to areas of pathogen entry. The heightened sensitivity of the ED zone was also reflected in our genome-wide transcriptomic results. Compared to the whole root samples, a higher number of DEGs correlating with plant-biotic interactions and PTI were also seen in ED samples.

Our results also show a tightly regulated transcriptional reprogramming by FLS2 and FLS3 with clear differences in number of upregulated DEGs for both flg22 (WR: 2145, ED: 2959) and flgII- 28 (WR: 248, ED: 1843) treatment. Although we found many overlapping genes between flg22 and flgII-28 treatments in the ED zone (1,496 genes) the response to flg22 had 1,453 distinct upregulated DEGs compared to 335 genes for flgII-28.

Together, our data suggests that PTI-driven ROS formation, MPK1/2/3 activation, and transcriptional reprogramming are primarily located in specific developmental regions of the tomato root. In the context of tomato, the PTI response initiated by FLS3 appears to be a more specific process compared to FLS2, reflecting evolutionary adaptation in PTI response.

## Conclusions

Our work identifies key differences in FLS2, FLS3, and CORE-mediated PTI pathways in tomato roots, highlighting the importance of studying PTI across a range of plant species to understand the diversity and evolution of plant immune systems. Understanding how PRR pathways diverge and the impact on downstream phenotypes in different species provides a foundation for developing targeted strategies to optimize PTI responses and broad-spectrum resistance in crops species.

## Acknowledgements

This work was supported by NSF Graduate Research Fellowship DGE-1842166 to RLK and USDA National Needs 2020-38420-30722 grant to AIP as well as funding from Purdue University Hatch #7002587. DMS was supported by USDA-NIFA 2021-67034-35049 and GC was supported by the National Institutes of Health 1R35GM136402. We thank Gregory Martin for providing the *Slrbohb* seeds and Christopher Staiger for the Arabidopsis seeds. We also thank members of the Iyer-Pascuzzi and Helm labs for critical reading of the manuscript.

## Funding

USDA-NIFA National Needs Fellowship, NSF-GRFP Fellowship

## Supporting Information

**Fig. S1. Reactive Oxygen Species (ROS) burst dynamics vary by PAMP type for tomato whole roots.**

**Fig. S2. LA0176 does not respond to csp22^Rsol^.**

**Fig. S3: Reactive Oxygen Species Burst varies both among cultivars and between PAMP type in the ED zone**

**Fig. S4. qPCR of virus-induced gene silencing (VIGS) constructs confirming reduced expression.**

**Fig. S5. Top 20 GO Biological Function categories represented by genes upregulated in response to PAMP treatments in tomato roots.**

**Fig. S6. Temporary root growth inhibition is observed in *Arabidopsis* seedlings for flg22^Pto^ treatment at 24 hpi, but not earlier.**

**Fig. S7. Determination of DPI concentration sufficient to fully inhibit H7996 ROS burst in response to flg22^Pto^.**

**Table S1. Primers and Constructs used in this study.**

**Table S2. Reactive Oxygen Species Burst is primarily found in the Early Differentiation Zone for additional cultivars of tomato.**

## Notes

### Competing Interest Statement

The authors have declared no competing interest.

