## Supplemental Figure 8 for "Tomato roots exhibit distinct, development-specific responses to bacterial-derived peptides"

### Slide 1
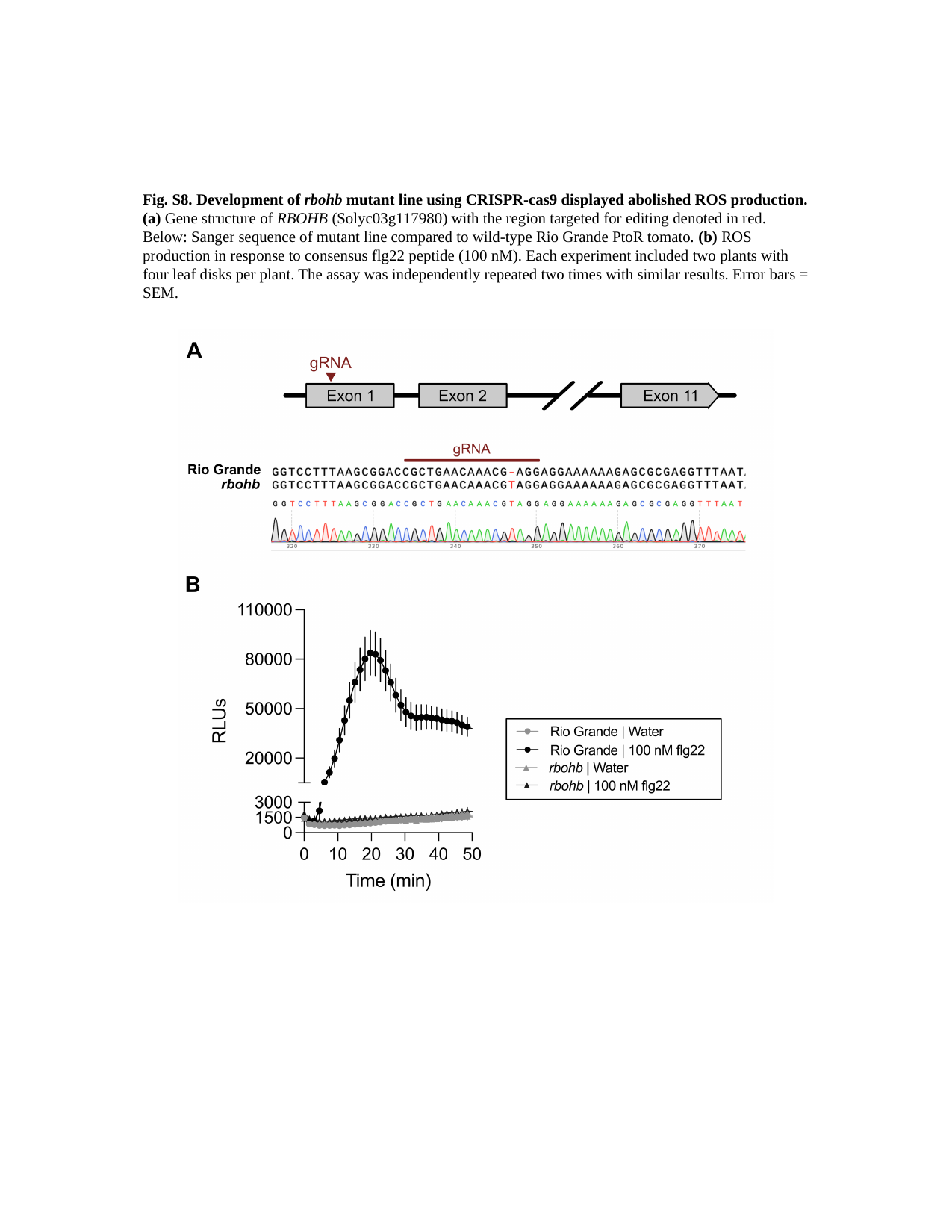

Fig. S8. Development of rbohb mutant line using CRISPR-cas9 displayed abolished ROS production. (a) Gene structure of RBOHB (Solyc03g117980) with the region targeted for editing denoted in red. Below: Sanger sequence of mutant line compared to wild-type Rio Grande PtoR tomato. (b) ROS production in response to consensus flg22 peptide (100 nM). Each experiment included two plants with four leaf disks per plant. The assay was independently repeated two times with similar results. Error bars = SEM.
