## Supplemental Figure 1 for "Tomato roots exhibit distinct, development-specific responses to bacterial-derived peptides"

### Slide 1
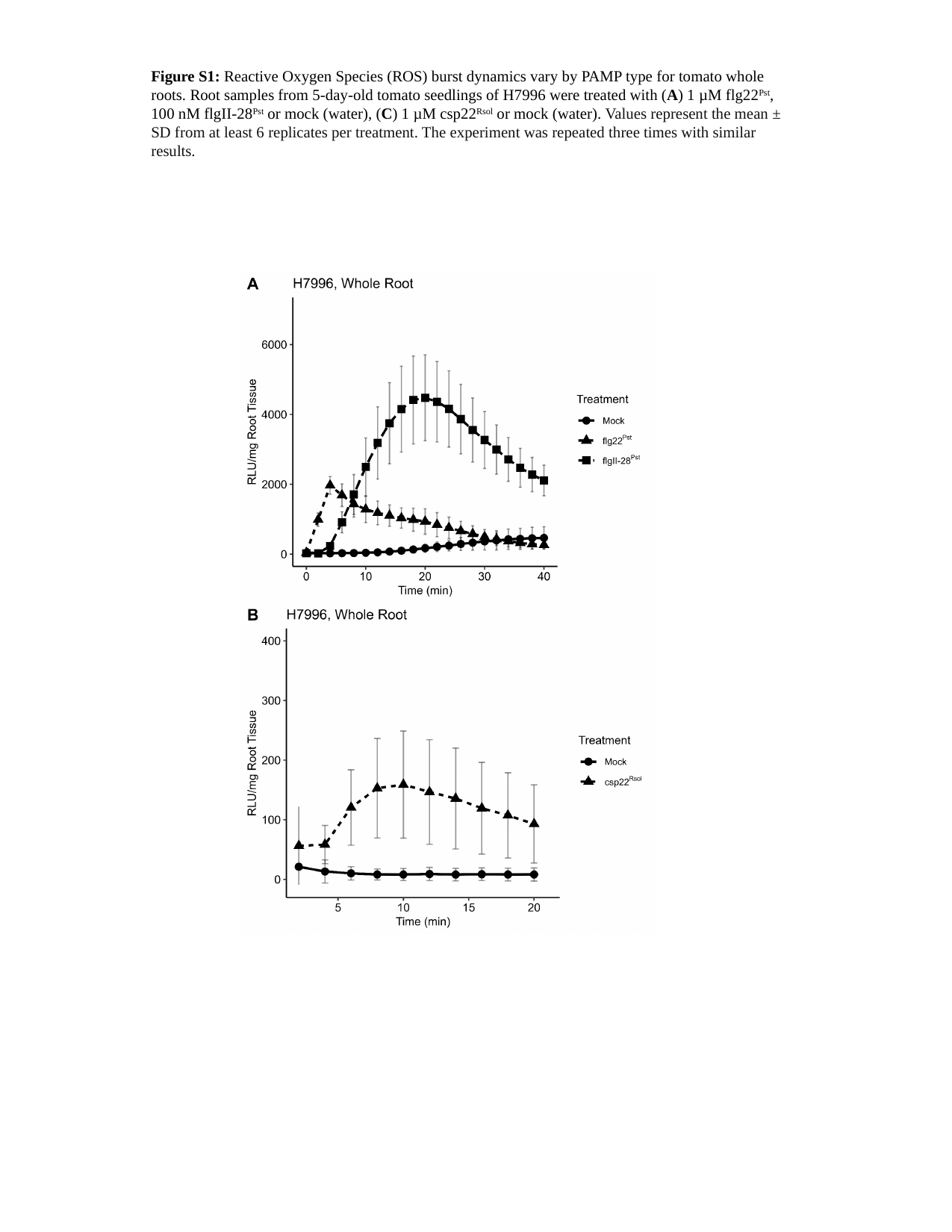

Figure S1: Reactive Oxygen Species (ROS) burst dynamics vary by PAMP type for tomato whole roots. Root samples from 5-day-old tomato seedlings of H7996 were treated with (A) 1 µM flg22^Pst^, 100 nM flgII-28^Pst^ or mock (water), (C) 1 µM csp22^Rsol^ or mock (water). Values represent the mean ± SD from at least 6 replicates per treatment. The experiment was repeated three times with similar results.
