## Supplemental Figure 2 for "Tomato roots exhibit distinct, development-specific responses to bacterial-derived peptides"

### Slide 1
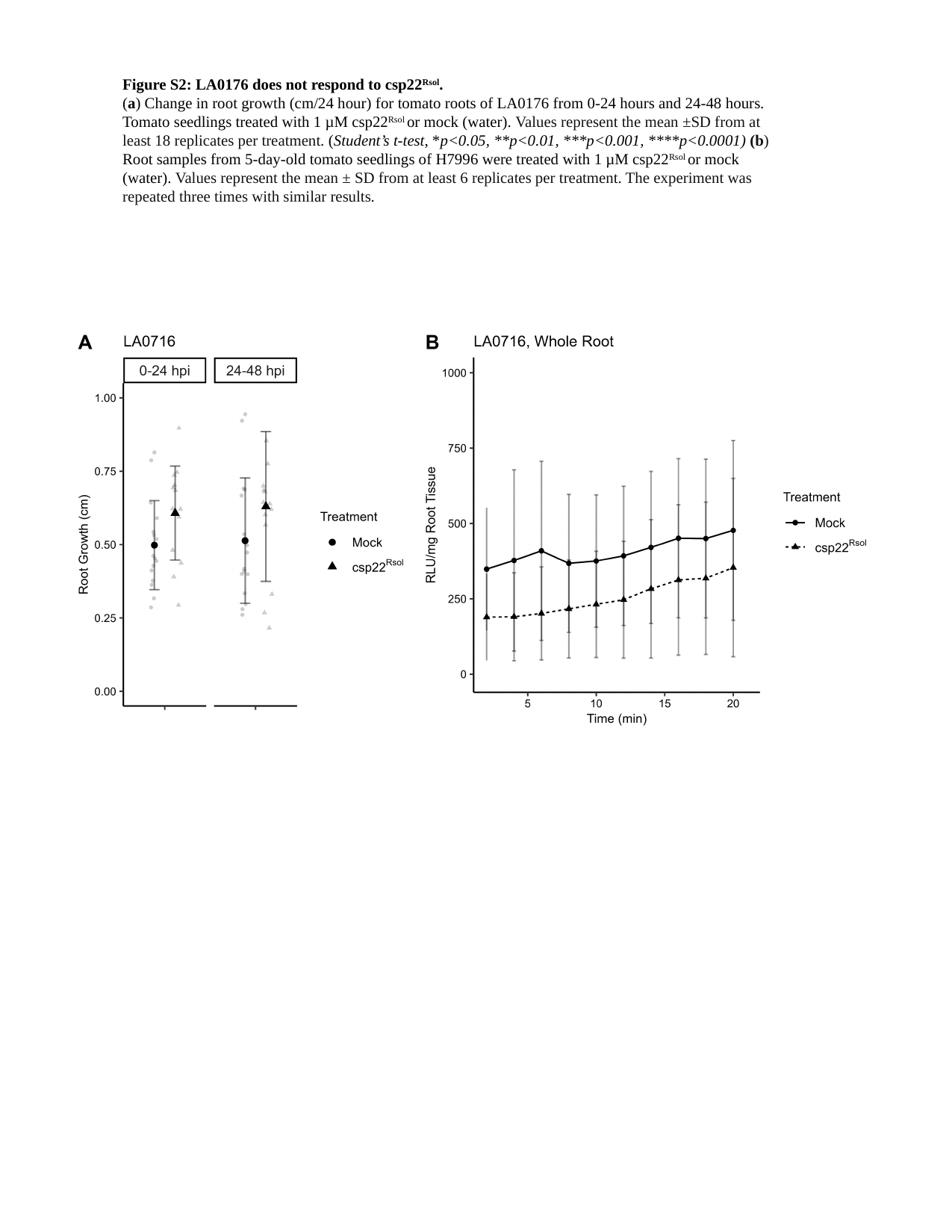

Figure S2: LA0176 does not respond to csp22^Rsol^.
(a) Change in root growth (cm/24 hour) for tomato roots of LA0176 from 0-24 hours and 24-48 hours. Tomato seedlings treated with 1 µM csp22^Rsol^ or mock (water). Values represent the mean ±SD from at least 18 replicates per treatment. (Student’s t-test, *p<0.05, **p<0.01, ***p<0.001, ****p<0.0001) (b) Root samples from 5-day-old tomato seedlings of H7996 were treated with 1 µM csp22^Rsol^ or mock (water). Values represent the mean ± SD from at least 6 replicates per treatment. The experiment was repeated three times with similar results.
