## Supplemental Figure 3 for "Tomato roots exhibit distinct, development-specific responses to bacterial-derived peptides"

### Slide 1
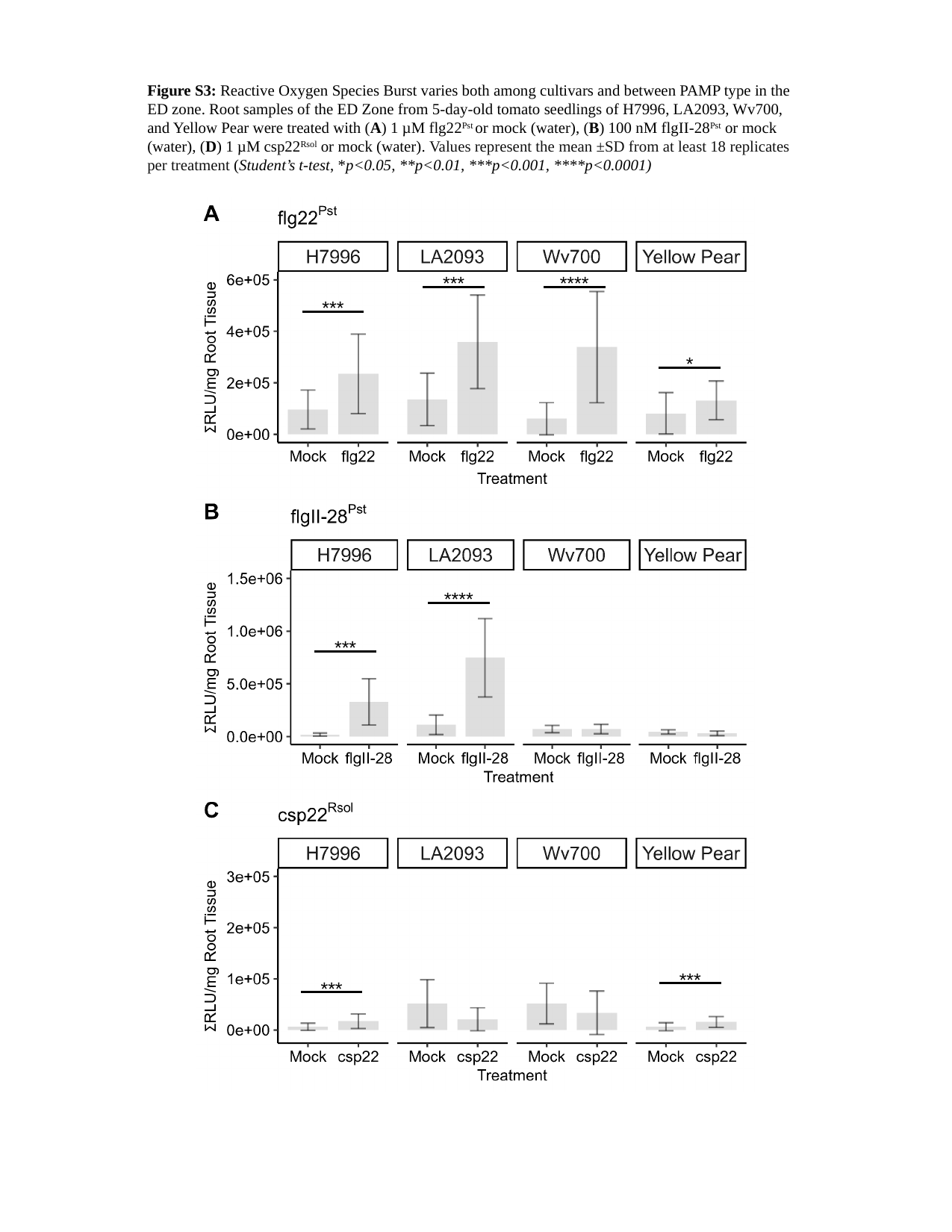

Figure S3: Reactive Oxygen Species Burst varies both among cultivars and between PAMP type in the ED zone. Root samples of the ED Zone from 5-day-old tomato seedlings of H7996, LA2093, Wv700, and Yellow Pear were treated with (A) 1 µM flg22^Pst^ or mock (water), (B) 100 nM flgII-28^Pst^ or mock (water), (D) 1 µM csp22^Rsol^ or mock (water). Values represent the mean ±SD from at least 18 replicates per treatment (Student’s t-test, *p<0.05, **p<0.01, ***p<0.001, ****p<0.0001)
***
****
***
*
****
***
***
***
