## Supplemental Figure 4 for "Tomato roots exhibit distinct, development-specific responses to bacterial-derived peptides"

### Slide 1
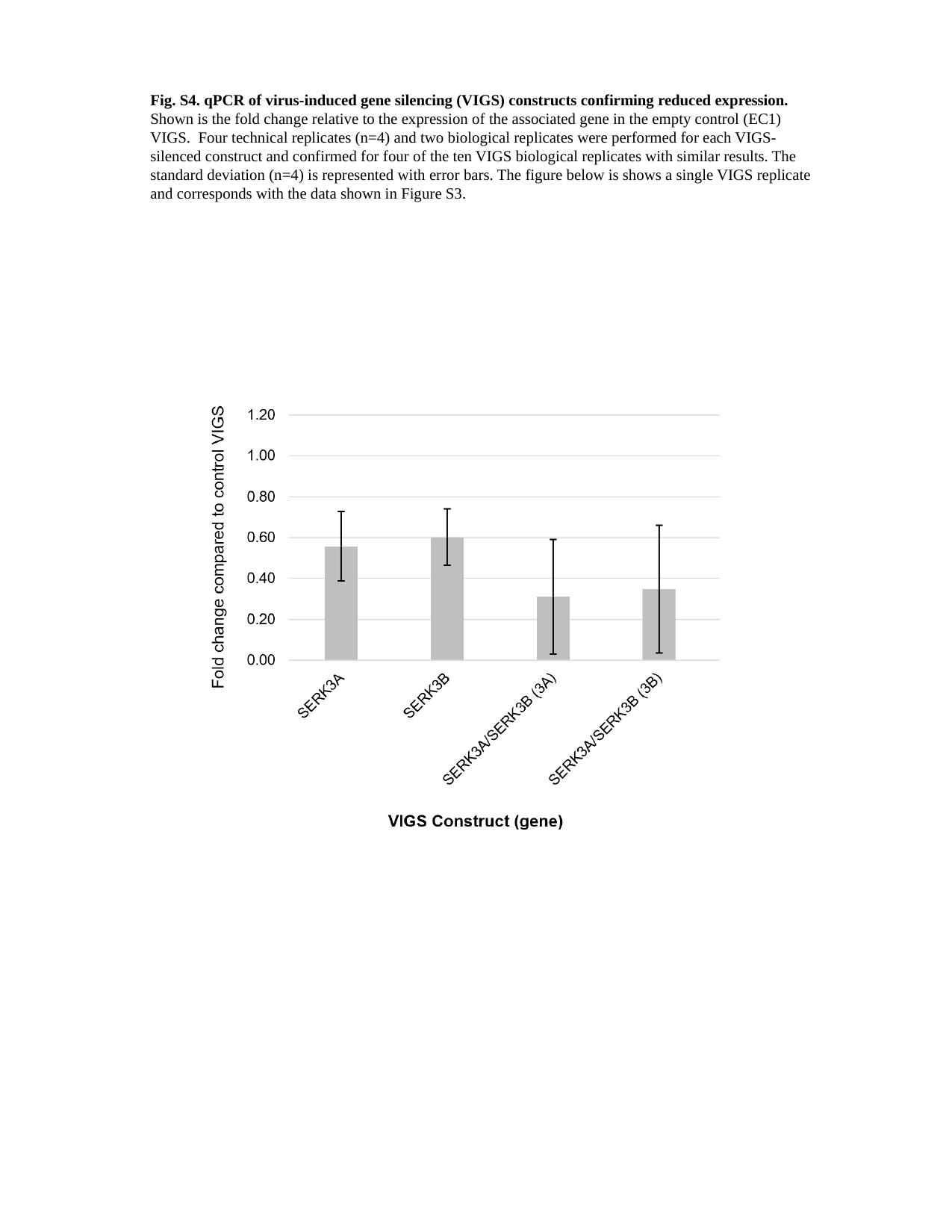

Fig. S4. qPCR of virus-induced gene silencing (VIGS) constructs confirming reduced expression. Shown is the fold change relative to the expression of the associated gene in the empty control (EC1) VIGS. Four technical replicates (n=4) and two biological replicates were performed for each VIGS-silenced construct and confirmed for four of the ten VIGS biological replicates with similar results. The standard deviation (n=4) is represented with error bars. The figure below is shows a single VIGS replicate and corresponds with the data shown in Figure S3.
