## Supplemental Figure 5 for "Tomato roots exhibit distinct, development-specific responses to bacterial-derived peptides"

### Slide 1
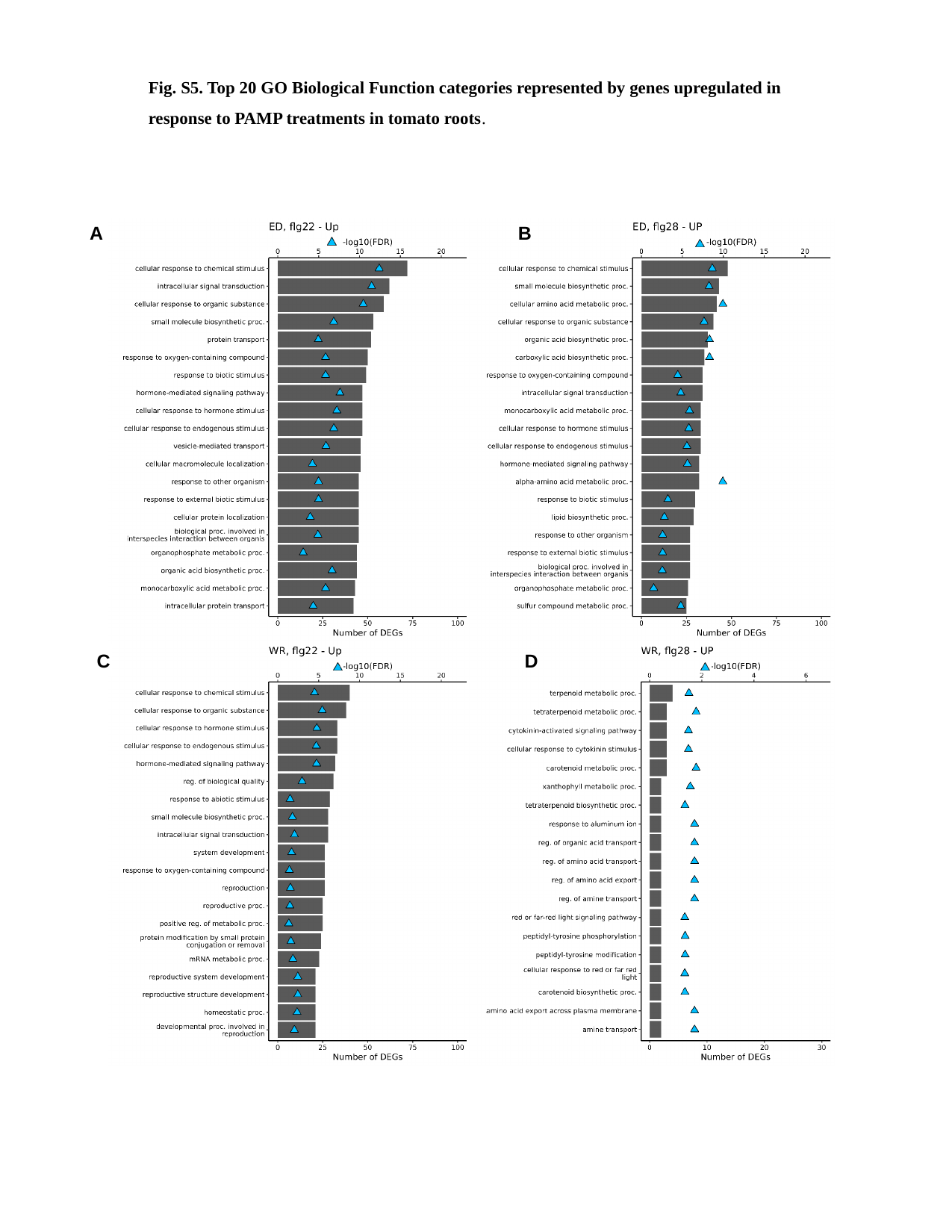

Fig. S5. Top 20 GO Biological Function categories represented by genes upregulated in response to PAMP treatments in tomato roots.
A
B
C
D
