## Supplemental Figure 6 for "Tomato roots exhibit distinct, development-specific responses to bacterial-derived peptides"

### Slide 1
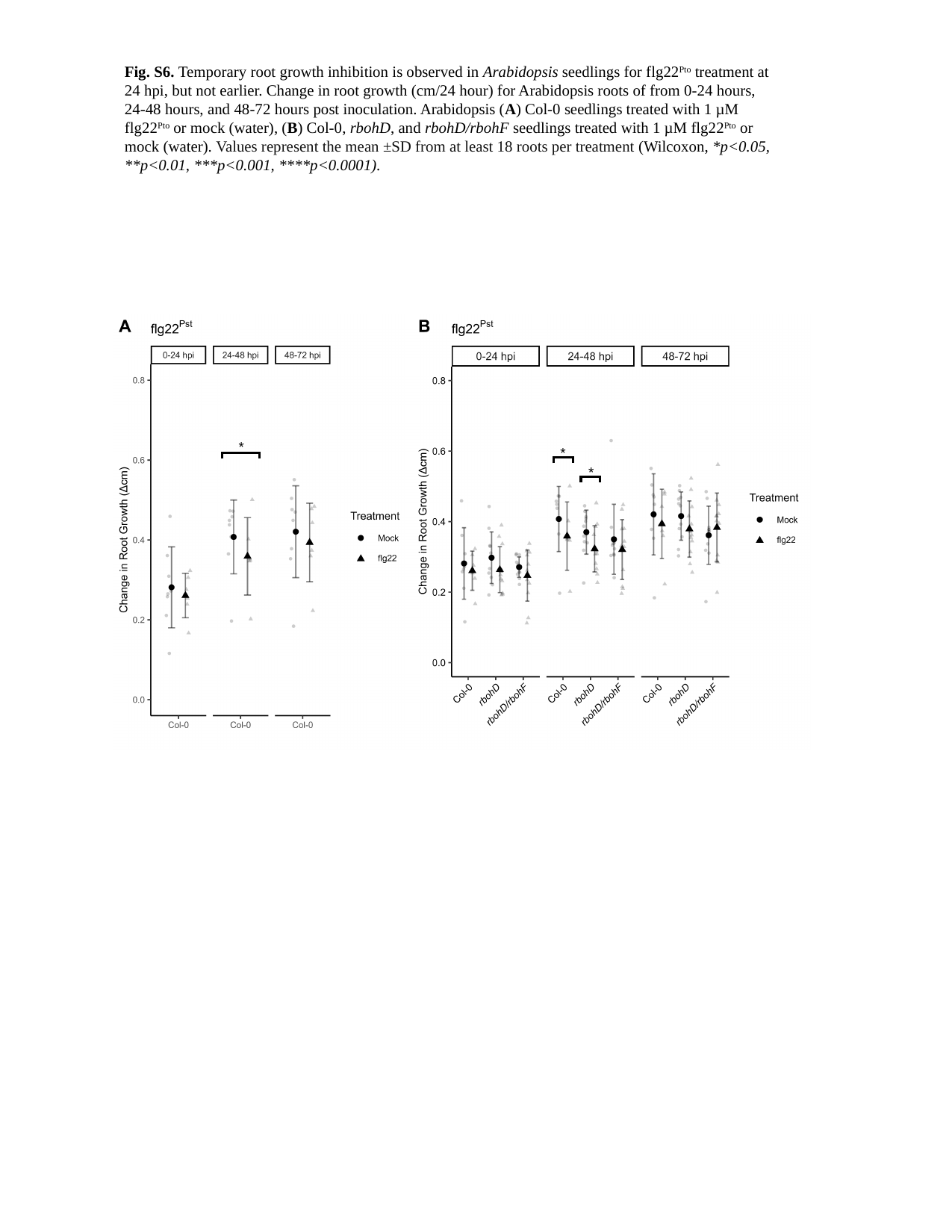

Fig. S6. Temporary root growth inhibition is observed in Arabidopsis seedlings for flg22^Pto^ treatment at 24 hpi, but not earlier.​ Change in root growth (cm/24 hour) for Arabidopsis roots of from 0-24 hours, 24-48 hours, and 48-72 hours post inoculation. Arabidopsis (A) Col-0 seedlings treated with 1 µM flg22^Pto^ or mock (water), (B) Col-0, rbohD, and rbohD/rbohF seedlings treated with 1 µM flg22^Pto^ or mock (water). Values represent the mean ±SD from at least 18 roots per treatment (Wilcoxon, *p<0.05, **p<0.01, ***p<0.001, ****p<0.0001).
*
*
*
