## Supplemental Figure 7 for "Tomato roots exhibit distinct, development-specific responses to bacterial-derived peptides"

### Slide 1
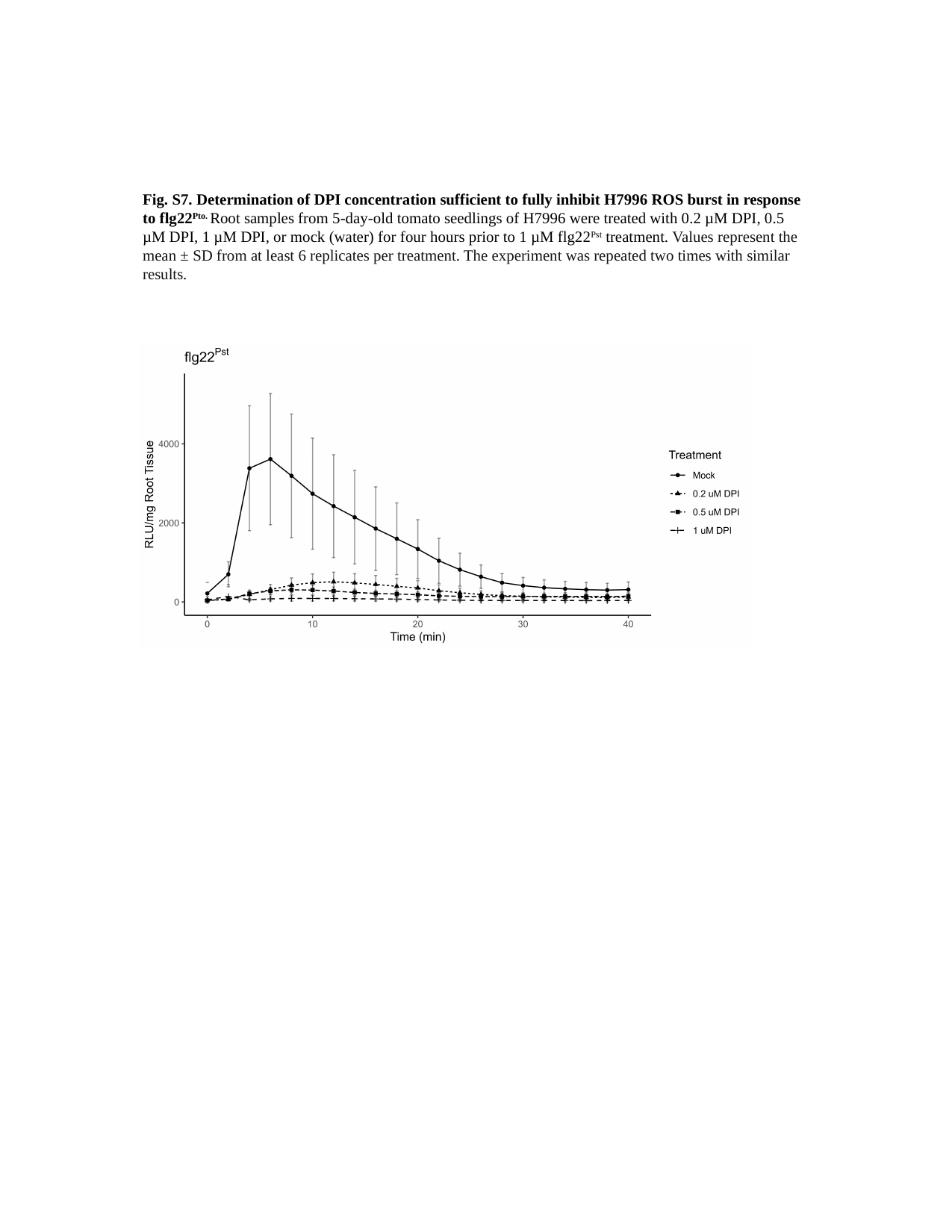

Fig. S7. Determination of DPI concentration sufficient to fully inhibit H7996 ROS burst in response to flg22^Pto.^ Root samples from 5-day-old tomato seedlings of H7996 were treated with 0.2 µM DPI, 0.5 µM DPI, 1 µM DPI, or mock (water) for four hours prior to 1 µM flg22^Pst^ treatment. Values represent the mean ± SD from at least 6 replicates per treatment. The experiment was repeated two times with similar results.
